# Epigenetic inheritance of centromere identity in a heterologous system

**DOI:** 10.1101/560391

**Authors:** Virginie Roure, Bethan Medina-Pritchard, Eduard Anselm, A. Arockia Jeyaprakash, Patrick Heun

## Abstract

The centromere is an essential chromosomal region required for accurate chromosome segregation. Most eukaryotic centromeres are defined epigenetically by the histone H3 variant, CENP-A, yet how its self-propagation is achieved remains poorly understood. Here we developed a heterologous system to reconstitute epigenetic inheritance of centromeric chromatin by ectopically targeting the *Drosophila* centromere proteins dCENP-A, dCENP-C and CAL1 to LacO arrays in human cells. Dissecting the function of these three components uncovers the key role of self-association of dCENP-C and CAL1 for their mutual interaction and dCENP-A deposition. Importantly, we identify the components required for dCENP-C loading onto chromatin, involving a cooperation between CAL1 and dCENP-A nucleosomes, thus closing the epigenetic loop to ensure dCENP-C and dCENP-A replenishment during the cell division cycle. Finally, we show that all three *Drosophila* factors are sufficient for dCENP-A propagation and propose a model for the epigenetic inheritance of centromere identity.

## INTRODUCTION

Centromeres are essential chromosomal regions that ensure the faithful distribution of genetic information during cell division. Most eukaryotic centromeres are defined by the presence of specialized nucleosomes containing the centromere-specific histone H3 variant, called CENP-A (Black and Cleveland, 2011). Consistent with the epigenetic nature of the centromere, CENP-A is found in every identified neocentromere (Marshall et al., 2008) and ectopic targeting of CENP-A to a non-centromeric chromosomal locus is sufficient to generate a functional and heritable centromere (Barnhart et al., 2011; Logsdon et al., 2015; Mendiburo et al., 2011). The key function of CENP-A is to provide a self-propagating platform that directs kinetochore assembly, a macromolecular structure which mediates chromosome attachment to spindle microtubules during mitosis and meiosis.

To ensure epigenetic inheritance the centromere mark must stably persist through the multiple division cycles of a cell. Unlike canonical histone H3, CENP-A nucleosomes are not replenished during DNA replication but from late mitosis through G1 phase (Dunleavy et al., 2012; Jansen et al., 2007; Lidsky et al., 2013; Schuh et al., 2007). One attractive model for the self-propagation of CENP-A is an epigenetic loop, where one or more adaptors recognize and direct the deposition of new CENP-A. In human, CENP-A (hCENP-A) deposition requires the coordinated activity of several factors: the hCENP-A specific histone chaperone HJURP, hCENP-C and the Mis18 complex (McKinley and Cheeseman, 2016). The members of the Mis18 complex have been shown to interact with hCENP-C (Dambacher et al., 2012; Moree et al., 2011) and to recruit HJURP to centromeres (Barnhart et al., 2011; Wang et al., 2014) to promote loading of hCENP-A. Direct binding of hCENP-C to hCENP-A nucleosomes (Carroll et al., 2010; Kato et al., 2013) closes the loop, thereby providing a mechanism to ensure CENP-A propagation in human centromeres.

In *Drosophila*, in addition to CENP-A (dCENP-A, also known as CID or cenH3), only two centromere proteins have been identified so far: the dCENP-A specific chaperone CAL1 that mediates dCENP-A deposition (Chen et al., 2014), and *Drosophila* CENP-C (dCENP-C) (Heeger et al., 2005). CAL1, dCENP-C and dCENP-A have been shown to be interdependent for centromere localization and function (Erhardt et al., 2008; Schittenhelm et al., 2010). However, in contrast to their human counterparts, dCENP-C and dCENP-A appear to interact only indirectly via the bridging factor CAL1, which binds dCENP-A through its N-terminal domain and dCENP-C through its C-terminal domain (Schittenhelm et al., 2010). CAL1 has been shown to be sufficient for dCENP-A nucleosome assembly and it has been proposed that dCENP-C mediates CAL1/dCENP-A recruitment to centromeres (Chen et al., 2014). However, how dCENP-C associates with the centromere and how centromeric chromatin is epigenetically propagated is not understood.

Although analysis of dCENP-A, dCENP-C and CAL1 in their ‘natural’ environment in *Drosophila* cells has provided insights into their roles in maintaining centromere identity, all three factors exhibit dependencies on each other for function and protein stability. The use of a heterologous system where none of the three proteins are essential for viability is unencumbered by these complexities. Hence to explore this possibility, we took advantage of the pronounced evolutionary divergence between *Drosophila* and human centromere components. Using the LacI/LacO system, we artificially targeted the three *Drosophila* centromere proteins dCENP-A, dCENP-C and CAL1 to chromosomally integrated LacO arrays in human U2OS cells to dissect their interactions and role in dCENP-A inheritance in unprecedented detail. First, we generated histone H3/dCENP-A chimeras to identify the *Drosophila* CENP-A centromere targeting domain as well as the interaction domain of dCENP-A with CAL1. LacI/LacO-targeting further revealed the joint roles of both CAL1 and dCENP-A in recruiting dCENP-C to chromatin and highlighted the importance of dCENP-C and CAL1 self-association for their interactions and dCENP-A deposition. Finally, we showed that these three factors are sufficient for propagation of dCENP-A and proposed a model for the epigenetic inheritance of centromere identity in *Drosophila*.

## RESULTS

### Identification of the centromere targeting domain of *Drosophila* CENP-A

To determine the region of *Drosophila* CENP-A required for its localization to centromeres, we designed a collection of chimeric dCENP-A/dH3 variants in which one or several domains of histone dH3 were replaced by the corresponding domains of histone dCENP-A. The secondary structure of the histone fold is composed of three helices (α1, α2, α3), which are connected by two loops (L1 and L2) (Figure 1A). Despite the divergence in amino-acid composition (overall ≈20%, histone fold ≈ 38% identity), dCENP-A mainly differs from dH3 in the longer loop 1 and N-terminal tail (Figures 1A and S1A). In human cells L1 and the α2-helix of hCENP-A are sufficient to target a H3-chimera to centromeres and are hence named CENP-A targeting domain (hCATD, Figure 1A) (Black et al., 2004). We divided *Drosophila* CENP-A and H3 into four regions: N-terminal (N), L1 loop, helix α2 and C-terminal part (C) and expressed variants of dCENP-A/dH3 chimera fused to the Hemagglutinin (HA)-tag in *Drosophila* Schneider S2 cells (Figures 1A-1C).

**Figure 1:**
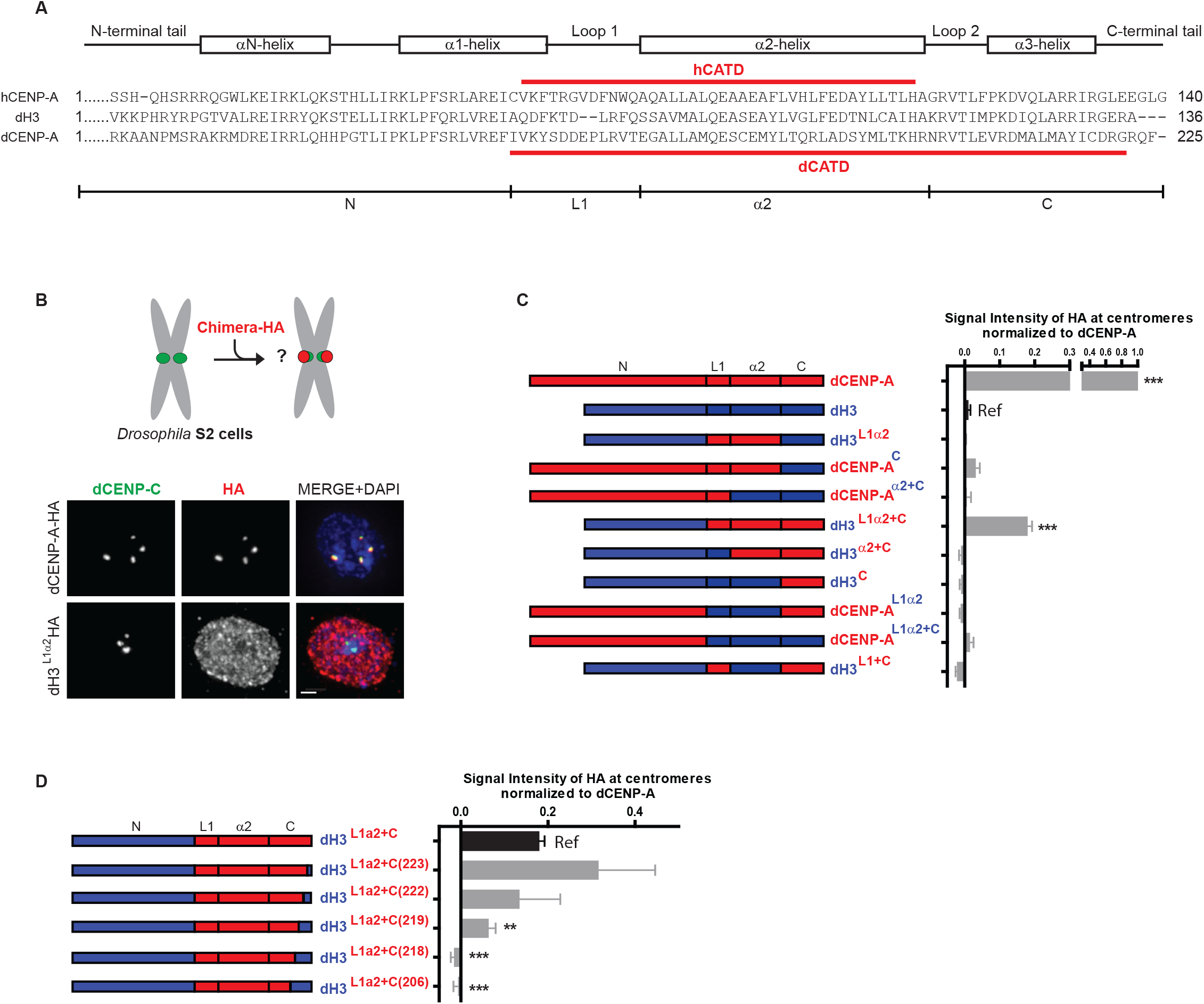
The CATD of CENP-A in *Drosophila* is larger than in humans and includes the a3 helix. (A) *Drosophila* CENP-A was divided into four domains: The N-terminal (N) from residues 1 to 160 (corresponding to residue 1 to 75 in dH3), the L1 domain from residue 161 to 173 contains loop L1 (residue 76 to 86 in dH3), the α2 domain contains helix α2 (residue 174 to 202 in dCENP-A, residue 87 to 115 in dH3) and the C-terminal (C) from residue 203 to 225 (residue 116 to 136 in dH3). (B) Experimental scheme and representative IF images of HA-tagged dCENP-A WT and the chimera dH3^L1α2^ expression patterns in S2 *Drosophila* cells. dCENP-C marks *Drosophila* centromeres. (C-D) Quantitation of indicated dCENP-A/dH3 chimera mean intensities at centromeres normalized to dCENP-A mean intensity at centromeres. Scale bar, 1μm. Error bars show SEM. Asterisks denote significant differences (** P < 0.01; *** P < 0.001), absence of asterisk denotes not significant difference. The reference sample for statistical analysis is indicated as Ref.

Targeting of each chimera to endogenous *Drosophila* centromeres was assessed by quantitative immunofluorescence (IF) where the intensity of signal was measured (Figures 1B and 1C). Surprisingly, we observed that, unlike in humans, the L1 and α2 regions of dCENP-A (dH3^L1α2^) are not sufficient and inclusion of the C region is required for dCENP-A/dH3 chimera to display centromere localization (dH3^L1α2+C^). However, the L1 and α2 regions are both necessary, given that omittance of any of these two regions (dH3^α2+C^ and dH3^L1+ C^) interferes with centromere targeting. Although the N region of dCENP-A is neither necessary nor sufficient for centromere targeting, it might still contribute to targeting, as the dH3^L1α2+C^ chimera is less efficient than wild type dCENP-A (around 20%).

Narrowing further down the domain required for centromere targeting showed that the α3-helix in the C-terminal region is absolutely essential and only the last three residues of dCENP-A can be omitted for centromere localization (Figure 1D). Taken together, the *Drosophila* centromere targeting domain comprises residues 161 to 222, extending the human CATD up to the α3-helix and is therefore significantly larger in dCENP-A (Figure 1A).

### Development of a heterologous system to reconstitute dCENP-A loading

The CATD of human CENP-A is also the domain bound by its chaperone HJURP (Bassett et al., 2012). Having defined the dCATD, we next sought to determine the domain of interaction between dCENP-A and its chaperone CAL1. To prevent interference due to the presence of endogenous centromere proteins in *Drosophila* cells, we developed a heterologous system in which *Drosophila* proteins were expressed in human cells.

Both experimental and bioinformatic approaches show little conservation between human and *Drosophila* for factors involved in CENP-A replenishment, revealing a low degree of sequence similarity for some components in *Drosophila* (CENP-A, CENP-C) and the complete absence of others (HJURP, Mis18 complex; Figure S1A)(McKinley and Cheeseman, 2016; Westermann and Schleiffer, 2013). Thus, we reasoned that human proteins would not interact or interfere with exogenously expressed *Drosophila* centromere proteins, yet providing both a similar chromatin context and physiological conditions to support protein interactions and function. Here, we utilized U2OS cells containing integrated mixed LacO and TetO arrays and transiently expressed *Drosophila* centromere proteins fused to the *lac* repressor (LacI) (Barnhart et al., 2011; Janicki et al., 2004). The LacI-fusion proteins can be specifically targeted to the LacO arrays and analyzed for the co-recruitment of other ectopically expressed *Drosophila* proteins of interest (Zolghadr et al., 2008).

To confirm that there is little or no crosstalk between the human and the *Drosophila* centromere proteins in these cells, we first expressed the *Drosophila* centromere proteins CAL1, dCENP-A and dCENP-C fused to a tandem GFP-LacI tag. Each GFP-LacI fusion was then targeted to the LacO arrays in U2OS cells and tested for their ability to recruit the human homologs (hCENP-A and hCENP-C) by IF with anti hCENP-A and anti hCENP-C specific antibodies (Figures S1B-1D). None of the human centromere proteins are recruited to the LacO arrays by *Drosophila* CENP-A, CENP-C or CAL1, similar to the GFP-LacI negative control (Figures S1C and S1D). As a positive control, the human centromere protein HJURP tagged with GFP-LacI significantly recruits hCENP-A (Figure S1C, lane1) and hCENP-C (Figure S1D, lane1) to the LacO arrays. Moreover, the absence of interactions between human and *Drosophila* centromere proteins was confirmed in the reciprocal heterologous system where human centromere proteins were expressed in *Drosophila melanogaster* S2 cells containing a LacO plasmid (Logsdon et al., 2015). We also noticed that dCENP-C was occasionally localized at human centromeres, but decided to exclusively focus on the LacO array, where we found no interference. We conclude there is no significant association of endogenous human centromere proteins with the exogenously expressed *Drosophila* centromere proteins at the LacO array in our heterologous approach.

### The dCATD is required for wildtype levels of CAL1/dCENP-A interaction

To determine the domain of interaction of dCENP-A with CAL1, dCENP-A/dH3 chimeras expressed as GFP-LacI fusions were tethered to LacO arrays in U2OS transiently expressing V5-tagged CAL1 to analyse its localization (Figure 2A). Although weak association between CAL1 and the dH3^L1α2^ chimera are detected, the presence of the N (dCENP-A^C^) or C part of dCENP-A (dH3^L1α2+C^, dH3^L1α2+C(222)^, dH3^L1α2+C(219)^, dH3^L1α2+C(206)^) significantly increases the co-recruitment of CAL1 to LacO arrays (Figure 2B). Generally, the level of interaction between the chimeras and CAL1 correlate with their ability to be targeted to *Drosophila* centromeres, with the full dCATD containing chimeras displaying the strongest recruitment of CAL1-V5 similar to wildtype dCENP-A (Figures 1C and 1D). However, the chimera dCENP-A^C^ exhibits lower but significant levels of CAL1 recruitment (60% compare to WT dCENP-A, Figure 2B), yet hardly localizes to centromeres (Figure 1C). This observation suggests that, although multiple regions across dCENP-A including its N region contribute to CAL1 binding, only the full dCATD provides the high level of association required for centromere targeting. In summary, these results show that the domain of interaction of dCENP-A with CAL1 is similar to the dCENP-A centromere targeting domain.

**Figure 2:**
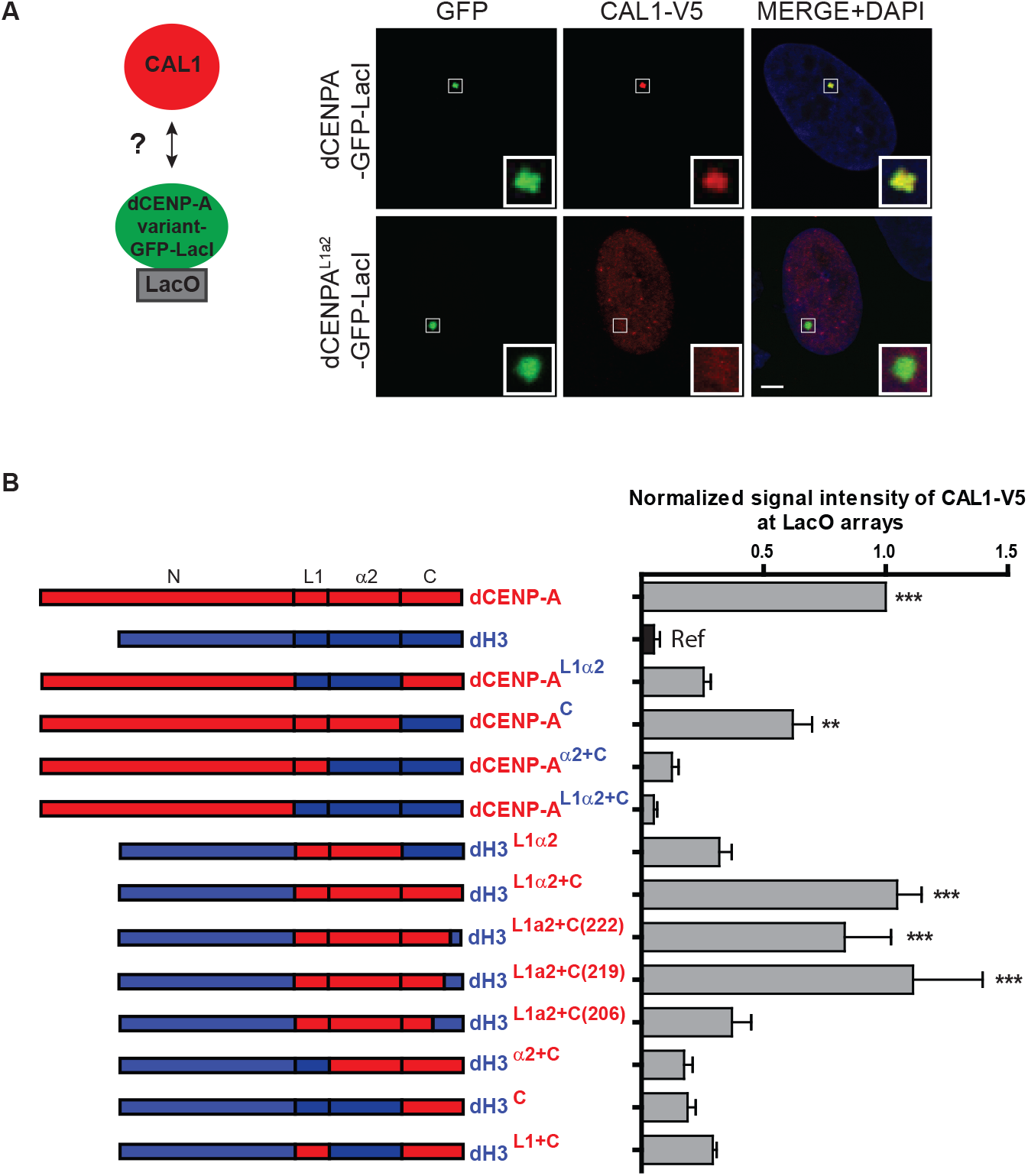
The domain of interaction of dCENP-A with CAL1 is similar to the dCATD domain. (A) Experimental scheme and representative IF images of CAL1-V5 recruitment to the LacO arrays by GFP-LacI-tagged dCENP-A and dCENP-A^L1α2^ in U2OS cells. (B) Quantitation of CAL1-V5 mean intensity at LacO upon tethering of the indicated GFP-LacI-tagged chimeras, normalized to its mean intensity at LacO upon tethering of GFP-LacI-tagged dCENP-A. Error bars show SEM. Asterisks denote significant differences (** P < 0.01; *** P < 0.001), absence of asterisk denotes not significant difference. Insets show magnification of the boxed regions. Scale bar, 5μm. The reference sample for statistical analysis is indicated as Ref.

### dCENP-C can recruit the CAL1/dCENP-A/H4 complex by direct interaction with CAL1

Taking advantage of our heterologous system to investigate direct interactions, we decided to further dissect step-by-step dCENP-A and CAL1 recruitment to chromatin. Initially we tested whether the previously reported direct interaction of dCENP-A and CAL1 can be reproduced (Chen et al., 2014; Schittenhelm et al., 2010), and then if the presence of dCENP-C could enhance the association of dCENP-A and CAL1. As expected, GFP-LacI tethered CAL1 can efficiently recruit dCENP-A to the LacO and vice versa, similarly in presence or absence of HA-dCENP-C, as compared to the GFP-LacI control (Figures 3A-3C and S2A), confirming previous findings that CAL1 and dCENP-A directly interact.

To investigate if dCENP-A nucleosomes maintain affinity for CAL1, we first checked whether dCENP-A-GFP-LacI is incorporated into chromatin assembled on LacO arrays. As previously shown with human CENP-A (Barnhart et al., 2011; Logsdon et al., 2015), we expected to find two pools of LacI-tagged dCENP-A at the LacO arrays: one bound to LacO through the LacI tether and sensitive to the LacI allosteric effector molecule IPTG; and another that is stably assembled into nucleosomes and therefore IPTG insensitive. dCENP-A was tagged with a mutated LacI variant (LacIw) with increased IPTG sensitivity (Loiodice et al., 2014) and co-transfected with HA-tagged CAL1 and V5-tagged Tet repressor (TetR) to independently mark the arrays. 48h hours after transfection, cells were treated with IPTG and analyzed to determine the IPTG-sensitivity of dCENP-A and CAL1. While we found that the control GFP-LacIw is almost entirely displaced (Figures 3D and 3E), roughly half of the pool of dCENP-A-GFP-LacIw is retained at the LacO arrays after IPTG treatment (Figures 3F and 3G), suggesting that a fraction of dCENP-A has assembled into chromatin. As a control for proper dCENP-A incorporation into human chromatin, we mutated two hydrophobic residues in dCENP-A, Y187A and L188Q predicted to disrupt its interaction with histone dH4 (equivalent to F101 and L102 in the hCENP-A crystal structure; (Sekulic et al., 2010) and prevent its chromatin loading. Indeed, this dCENP-A mutant is completely removed from chromatin upon addition of IPTG (Figure S2B), suggesting that wildtype dCENP-A can form a nucleosome with human H4 and be properly incorporated into chromatin. Indeed, gel filtration and SEC-MALS experiments confirm that CAL1 forms a complex with recombinant dCENP-A and human histone H4 (Figures S2C and S2D). Importantly, the co-recruitment of CAL1-HA to the LacO arrays by dCENP-A-GFP-LacIw is completely IPTG-sensitive (Figures 3F and 3G), indicating that CAL1 cannot bind nucleosomal dCENP-A and that only the pool of non-nucleosomal dCENP-A is available for binding its chaperone CAL1, presumably mimicking the soluble form of the protein. Together, these results strongly suggest that once recruited to centromeres and after chromatin loading of dCENP-A, CAL1 behaves similar to human HJURP and dissociates from nucleosomal dCENP-A. Importantly, it also shows that nucleosomal dCENP-A does not directly recruit a new CAL1/dCENP-A/H4 soluble complex.

**Figure 3:**
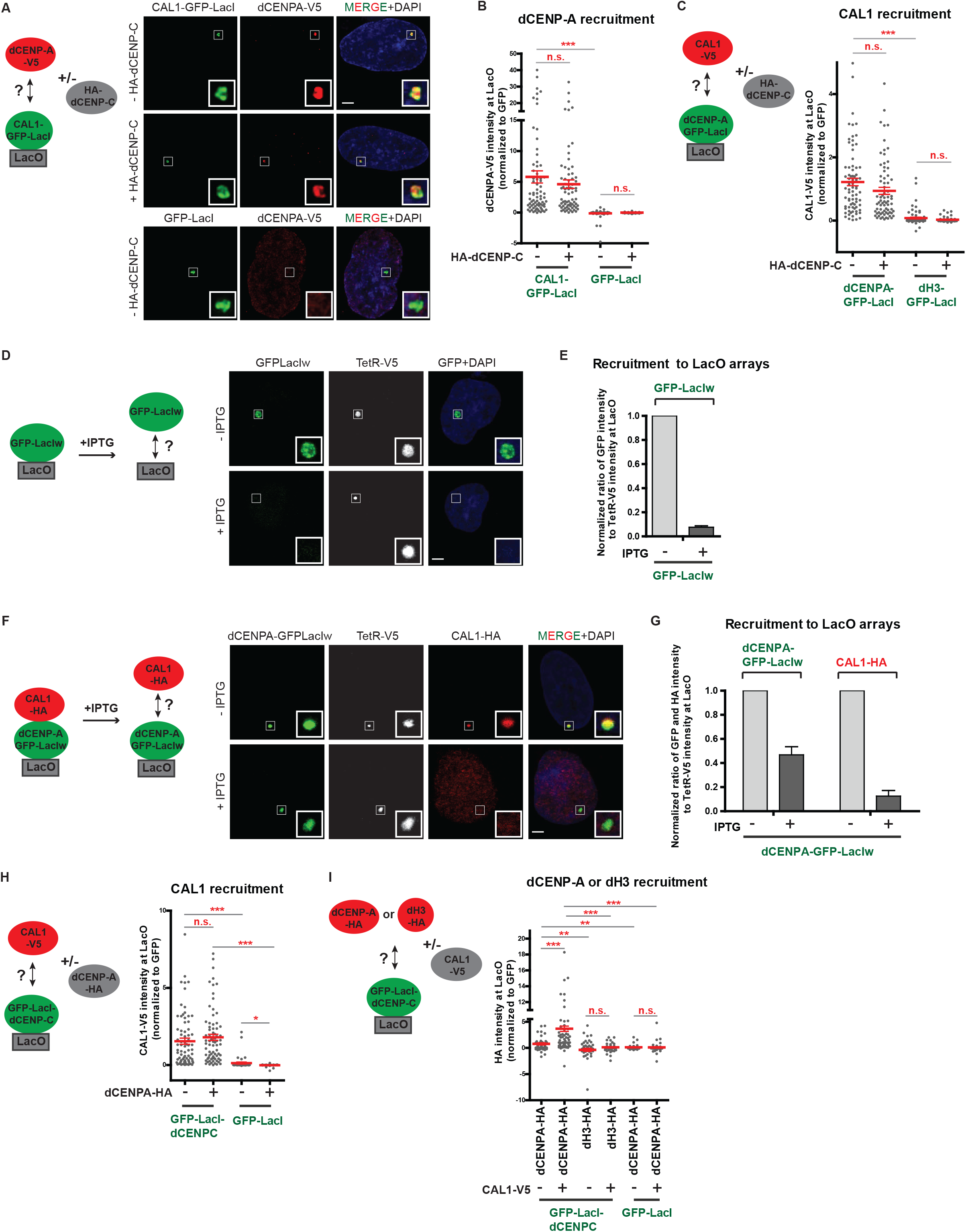
dCENP-C is the factor recruiting the complex CAL1/dCENP-A to the chromatin. (A) Experimental scheme and representative IF images of dCENP-A-V5 recruitment to the LacO arrays by CAL1-GFP-LacI or GFP-LacI, +/− HA-dCENP-C, in U2OS cells. (B) Quantitation of normalized dCENP-A-V5 mean intensity at LacO upon tethering of CAL1-GFP-LacI or the control GFP-LacI, +/− HA-dCENP-C. (C) Experimental scheme and quantitation of normalized CAL1-V5 mean intensity at LacO upon tethering of GFP-LacI-tagged dCENP-A or dH3, +/− HA-dCENP-C. (D) Experimental scheme and representative IF images of GFP-LacIw recruitment to the LacO arrays, +/− IPTG treatment in U2OS cells. (E) Quantitation of normalized GFP-LacIw mean intensity at LacO, +/− IPTG treatment. (F) Experimental scheme and representative IF images of CAL1-HA recruitment to the LacO arrays by dCENP-A-GFP-LacIw, +/− IPTG treatment in U2OS cells. (G) Quantitation of normalized dCENP-A-GFP-LacIw and CAL1-HA mean intensities at LacO, +/− IPTG treatment. (H) Experimental scheme and quantitation of normalized CAL1-V5 mean intensity at LacO upon tethering of GFP-Lac-dCENP-C or GFP-LacI, +/− dCENP-A-HA. (I) Experimental scheme and quantitation of normalized HA-tagged dCENP-A or dH3 mean intensities at LacO upon tethering of GFP-Lac-dCENP-C or GFP-LacI, +/− CAL1-V5. (See also Figure S2). Scale bar, 5μm. Insets show magnification of the boxed regions. Error bars show SEM (*P < 0.05; ** P < 0.01; *** P < 0.001; (n.s.) as not significant).

We next investigated whether dCENP-C is able to directly recruit CAL1, dCENP-A or both to the chromatin. We observed that, when GFP-LacI-dCENP-C is tethered to the LacO, it can recruit CAL1-V5 very efficiently in presence or absence of dCENP-A-HA, compared to the GFP-LacI negative control (Figures 3H and S2E). Surprisingly, we also observed low but significant direct interactions between GFP-LacI-dCENP-C and dCENP-A-HA in absence of CAL1 (Figures 3I and S2F), similar to direct interactions described between hCENP-C and hCENP-A (Carroll et al., 2010). However, the recruitment of dCENP-A is strongly enhanced by CAL1-V5 (Figures 3I and S2F). Hence this setup mimics the first step in dCENP-A loading and suggests that recognition of dCENP-C by CAL1 is required to recruit the CAL1/dCENP-A/dH4 soluble complex for the deposition of dCENP-A.

### CAL1 self-association is not required for binding but for deposition of dCENP-A

Dimerization of HJURP has been shown to be essential for the deposition of hCENP-A nucleosomes (Zasadzinska et al., 2013). We first checked if the property of self-association was conserved in the *Drosophila* histone chaperone CAL1. We observed that CAL1-GFP-LacI is able to recruit CAL1-HA to the arrays, suggesting that CAL1 can self-associate (Figure 4B, first and second lane; Figure S3A). To determine the CAL1 self-association domain, we generated different CAL1 fragments corresponding to the N-terminal (1-407), middle part (392-722) and C-terminal (699-979) of CAL1 as previously described (Schittenhelm et al., 2010)(Figure 4A) and performed the above described LacI/LacO recruitment assay. We found that only the fragment (1-407) is recruited by CAL1-GFP-LacI to the LacO arrays (Figures 4B and S3A), suggesting its ability to self-associate. By generating two fragments from residues 1 to 160 and from 101 to 407 (Chen et al., 2014)(Figure 4A), the CAL1 self-association domain was further narrowed down to the region contained between residues 161 and 407 of the N-terminal part (Figures 4B and S3A). SEC-MALS analysis confirms that the CAL1(1-160) fragment behaves as a monomer (Figure S3B).

**Figure 4:**
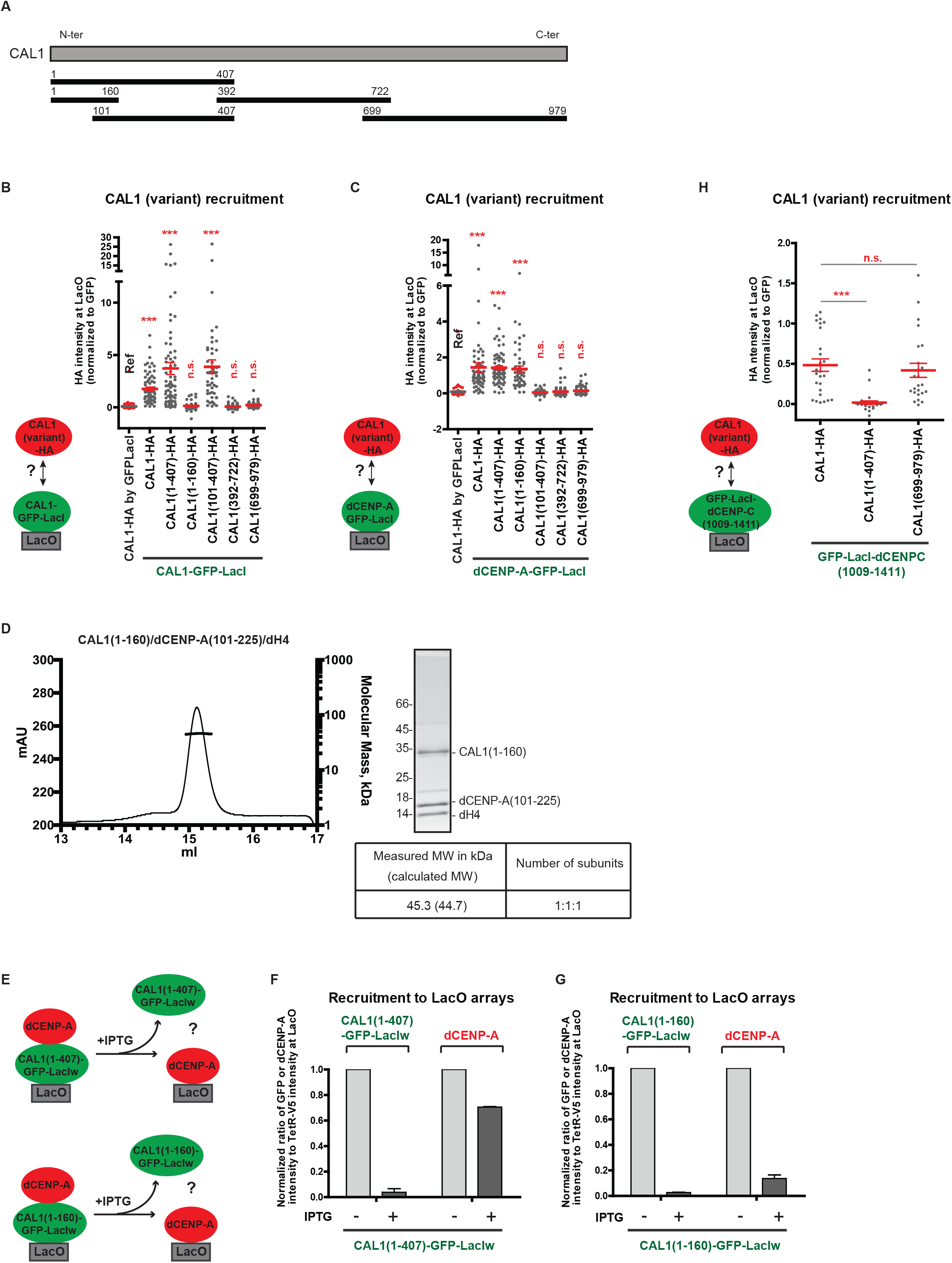
Self-association of CAL1 is not required for its interaction with the other centromere factors but for its chaperone activity and its centromere targeting. (A) The CAL1 N-terminal, middle and C-terminal regions used in this study are indicated by black horizontal lines with numbers indicating amino-acid positions. (B, C) Experimental schemes and quantitation of normalized HA-tagged CAL1 and CAL1 fragment mean intensities at LacO upon tethering of CAL1-GFP-LacI (B), dCENP-A-GFP-LacI (C) or GFP-LacI (B, C, first lane). (D) SEC-MALS experiments performed with recombinant CAL1(1-160) in complex with dCENP-A(101-225) and dH4. Elution volume (ml, x-axis) is plotted against absorption at 280 nm (mAu, left y-axis) and molecular mass (kDa, right y-axis). Tables show measure molecular weight (MW) and calculated MW. Mw/Mn = 1.001. (E) Experimental scheme for the experiments shown in (F, G). (F, G) Quantitation of normalized dCENP-A, CAL1(1-407)-GFP-LacIw (F) and CAL1(1-160)-GFP-LacIw (G) mean intensities at LacO, +/− IPTG treatment. (H) Quantitation of normalized HA-tagged CAL1 and CAL1 fragment mean intensities at LacO upon tethering of GFP-LacI-CENP-C(1009-1411). (See also Figures S3 and S4). Error bars show SEM (*** P < 0.001; (n.s.) as not significant). The reference sample for statistical analysis is indicated as Ref.

As CAL1(1-407) was previously shown to interact with dCENP-A (Schittenhelm et al., 2010), we asked if CAL1 self-association was required for its interaction with dCENP-A. CENP-A-GFP-LacI was tethered to the lacO arrays to determine the recruitment of the different CAL1-HA fragments. We first confirmed that the interaction domain of CAL1 with dCENP-A is contained between residues 1 to 160 of its N-terminal part (Figures 4C and S3C). Indeed, this result is consistent with previous *in vitro* experiments showing that the 1-160 fragment of CAL1 forms a complex with dCENP-A and dH4 (Chen et al., 2014) and was also confirmed by performing SEC-MALS experiments (Figure 4D). Together, these results suggest that different regions of CAL1 are responsible for its self-association and interaction with dCENP-A, suggesting that they are functionally separate.

To investigate further the role of CAL1 self-association in dCENP-A deposition, we next sought to understand the stochiometry of the CAL1/dCENP-A complex by performing SEC-MALS experiments in presence of CAL1(1-160), dCENP-A (101-255; lacking the N-terminal tail) and dH4 (Figure 4D). Interestingly, CAL1(1-160) binds only one (dCENP-A/dH4) dimer *in vitro*, suggesting that the self-association domain could be necessary to bring together two (dCENP-A/dH4) dimers to form a tetramer that is then deposited to the chromatin. To test this hypothesis, we determined whether CAL1 self-association was required for dCENP-A assembly into chromatin *in vivo* by analysing the persistence of dCENP-A at the LacO arrays after IPTG treatment upon tethering of GFP-LacIw-fusion of CAL1(1-407) or CAL1(1-160) (Figures 4E-4G). As expected, dCENP-A recruited by CAL1(1-407)-GFP-LacIw is not displaced from the arrays after IPTG treatment (Figures 4F and S3D), showing that dCENP-A is properly incorporated into chromatin by CAL1(1-407). Importantly, dCENP-A recruited by CAL1(1-160)-GFP-LacIw is IPTG-sensitive (Figures 4G and S3E), demonstrating that CAL1(1-160) missing its self-association domain is not able to incorporate dCENP-A into chromatin. This is in contrast to *in vitro* experiments, demonstrating that CAL1(1-160) is sufficient to assemble CENP-A nucleosomes using a plasmid supercoiling assay (Chen et al., 2014). In summary, our results strongly suggest that CAL1 self-association is necessary for deposition of the tetramer dCENP-A/dH4 to the chromatin. Thus, self association of both CAL1 and HJURP is critical for CENP-A deposition in *Drosophila* and human cells, respectively.

### Self-association of CAL1 is required for centromere targeting but not for interaction with dCENP-C

We next considered that CAL1 self-association might also be important for its interaction with dCENP-C. It has previously been shown that CAL1 and dCENP-C interact through their C-terminal domains (Schittenhelm et al., 2010). The C-terminal fragment of dCENP-C (residues 1009-1411) tethered to LacO arrays was analyzed for its ability to recruit full length CAL1 (containing the self-association domain), CAL1(699-979) (no self-association) or the negative control CAL1(1-407) (self-associating, but not interacting with dCENP-C). We observe that dCENP-C(1009-1411) can interact with full length CAL1 and CAL1 (699-979) with similar efficiencies, suggesting that CAL1 self-association is not required for its interaction with dCENP-C (Figures 4H and S4A).

However, we noticed that full length dCENP-C more effectively recruits full length CAL1 than CAL1(699-979), suggesting that the N-terminus of CAL1 could contribute to dCENP-C binding (Figure S4B), possibly due to its ability to self-associate. Expressing the (1-407) or (699-979) regions of CAL1 in S2 cells shows that neither is able to localize to *Drosophila* centromeres (Figures S4C and S4D), consistent with previous findings showing that CAL1 centromere localization requires the presence of its N- and C-terminal regions and depends on the interaction with both dCENP-A and dCENP-C. Moreover, dCENP-A and dCENP-C have been shown to be required for normal centromere localization of CAL1 by several studies (Erhardt et al., 2008; Goshima et al., 2007; Schittenhelm et al., 2010). In summary, this data suggests that CAL1 is recruited by dCENP-C as a multimer, possibly a dimer in complex with two dCENP-A/dH4.

### CAL1 efficiently recruits dCENP-C only in the presence of nucleosomal dCENP-A

One aspect of the epigenetic self-propagation loop of dCENP-A that remains poorly understood is how dCENP-C is recruited to the centromeres. To address this question, we first analyzed the recruitment of dCENP-C by dCENP-A, in presence or absence of CAL1. We found that tethered dCENP-A-GFP-LacI can recruit low levels of HA-dCENP-C in absence of CAL1-V5 (as compared to the H3-GFP-LacI control), (Figures 5A and 5B), confirming direct interaction of dCENP-A with dCENP-C (Figure 3I). The presence of CAL1 only slightly increases dCENP-C recruitment by dCENP-A-GFP-LacI (Figures 5A and 5B), possibly because CAL1 and dCENP-C do not appear to form a soluble complex (Mellone et al., 2011). Thus, dCENP-A alone is unlikely the only dCENP-C recruitment factor.

**Figure 5:**
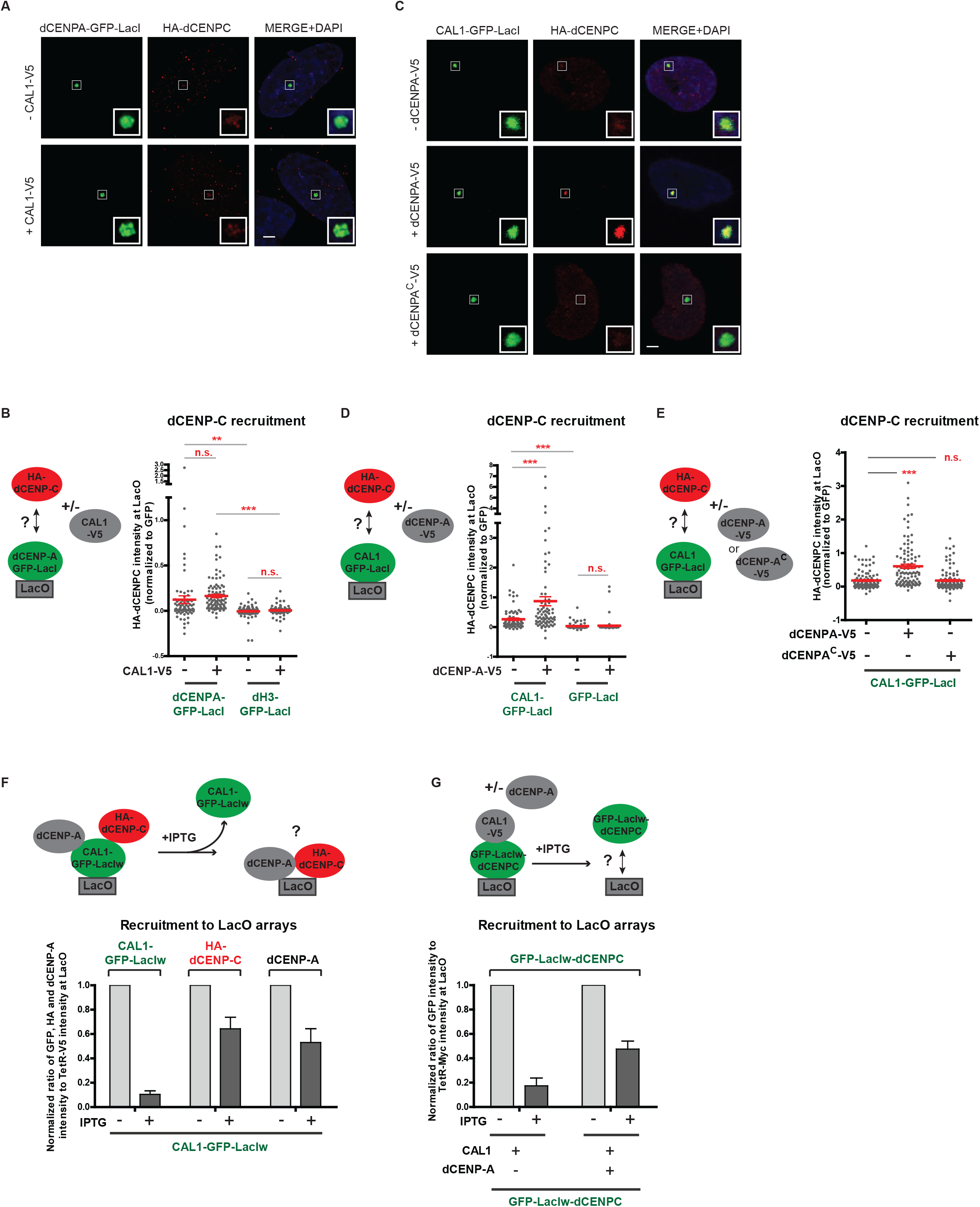
dCENP-C is loaded by CAL1 to the lacO arrays containing dCENP-A nucleosomes. (A) Representative IF images of HA-dCENP-C recruitment to the LacO arrays by dCENP-A-GFP-LacI, +/− CAL1-V5, in U2OS cells. (B) Experimental scheme and quantitation of normalized HA-dCENP-C mean intensity at LacO upon tethering of GFP-LacI-tagged dCENP-A-or dH3, +/− CAL1-V5. (C) Representative IF images of HA-dCENP-C recruitment to the LacO arrays by CAL1-GFP-LacI, +/− dCENP-A-V5 or dCENP-A-V5, in U2OS cells. (D) Experimental scheme and quantitation of normalized HA-dCENP-C mean intensity at LacO upon tethering of CAL1-GFP-LacI or GFP-LacI, +/− dCENP-A-V5. (E) Experimental scheme and quantitation of normalized HA-dCENP-C mean intensity at LacO upon tethering of CAL1-GFP-LacI, +/− V5-tagged dCENP-A or dCENP-A^C^. (F) Experimental scheme and quantitation of normalized CAL1-GFP-LacIw, HA-CENP-C and dCENP-A mean intensities at LacO, +/− IPTG treatment. (G) Experimental scheme and quantitation of normalized GFP-LacIw-dCENP-C mean intensity at LacO in presence of CAL1-V5, +/− dCENP-A and +/− IPTG treatment. (See also Figure S5). Scale bar, 5μm. Insets show magnification of the boxed regions. Error bars show SEM (** P < 0.01; *** P < 0.001; (n.s.) as not significant).

We next determined the ability of CAL1 to recruit dCENP-C. Based on the reported interactions between dCENP-C and CAL1 in yeast two-hybrid assays (Schittenhelm et al., 2010), we were surprised to find that CAL1-GFP-LacI alone recruits only low levels of HA-dCENP-C to the LacO (Figures 5C and 5D). However, HA-dCENP-C localization to the LacO is significantly enhanced in the presence of dCENP-A-V5, indicating that CAL1 and dCENP-A cooperate for dCENP-C recruitment (Figures 5C and 5D). The GFP-LacI tag on the C-terminal of CAL1 does not prevent its association with dCENP-C, as N-terminally GFP-LacI-tagged CAL1 presents identical level of interaction with dCENP-C (Figure S5A). Focusing on the role of dCENP-A, we investigated if nucleosomal or soluble dCENP-A in complex with CAL1 is important for dCENP-C recruitment. We first took advantage of the dCENP-A^C^ chimera, which interacts with CAL1 (Figure 2B), but cannot be stably incorporated into the chromatin, as shown by IPTG treatment experiments (Figure S2B). The ability of tethered CAL1-GFP-LacI to recruit HA-dCENP-C alone was tested in presence of dCENP-A-V5 or the chimera dCENP-A^C^-V5 (Figures 5C, 5E and S5B). The presence of dCENP-A-V5, but not dCENP-A^C^-V5 increases significantly the association of HA-dCENP-C with the LacO arrays relative to HA-dCENP-C alone. This suggests that dCENP-C has some affinity for dCENP-A nucleosomes, which might contribute to either recruitment or stable chromatin binding of dCENP-C on chromatin. To test this hypothesis, we first determined whether HA-dCENP-C recruitment by CAL1-GFP-LacIw was IPTG-sensitive in the presence of dCENP-A (Figures 5F and S5C). As expected, CAL1-GFP-LacIw is almost completely displaced from the LacO following IPTG addition, whereas more than 50% of dCENP-A pool is retained. Surprisingly, more than half of the HA-dCENP-C pool also remains at the LacO array, suggesting that dCENP-C has affinity for dCENP-A chromatin.

Tethering of GFP-LacIw-CENP-C in cells expressing CAL1-V5 in the presence or absence of dCENP-A revealed that dCENP-C is significantly retained after IPTG treatment only in the presence of dCENP-A (Figures 5G, S5D and S5E). This suggests that assembled dCENP-A nucleosomes allow retention of GFP-LacIw-dCENP-C following IPTG addition and does not require continuous presence of CAL1. Our observation that low levels of direct interactions between dCENP-C and dCENP-A occurs in absence of CAL1 further supports this finding (Figures 3I and 5B). In summary, this data suggests that CAL1 in collaboration with dCENP-A is required to initially recruit dCENP-C, but only nucleosomal dCENP-A is required to maintain stable binding of dCENP-C to chromatin.

### dCENP-C dimerization is required for CAL1 association

It has been previously shown that the C-terminal region of CENP-C is required for its dimerization in budding yeast (Mif2p) and human (Cohen et al., 2008; Sugimoto et al., 1997). To determine if dimerization is conserved in *Drosophila* CENP-C, GFP-LacI-dCENP-C was tested for its ability to recruit transiently expressed full length HA-dCENP-C. Significant HA-dCENP-C recruitment, compared to the GFP-LacI control, suggests that CENP-C in *Drosophila* is able to self-associate (Figure 6B, first and second lane). Different HA-tagged dCENP-C fragments corresponding to the N-terminus (1-575), middle part (558-1038) and C-terminus (1009-1411) of dCENP-C as previously published (Heeger et al., 2005)(Figure 6A) revealed that the C-terminal region of dCENP-C is sufficient for dCENP-C self-association (Figures 6B and S6A). Further dissection narrowed down the minimal self-association domain down to residues 1263-1411 (Figures 6A, 6B and S6A) and SEC-MALS analysis of recombinant dCENP-C(1264-1411) showed dimer formation of this domain (Figure 6C).

**Figure 6:**
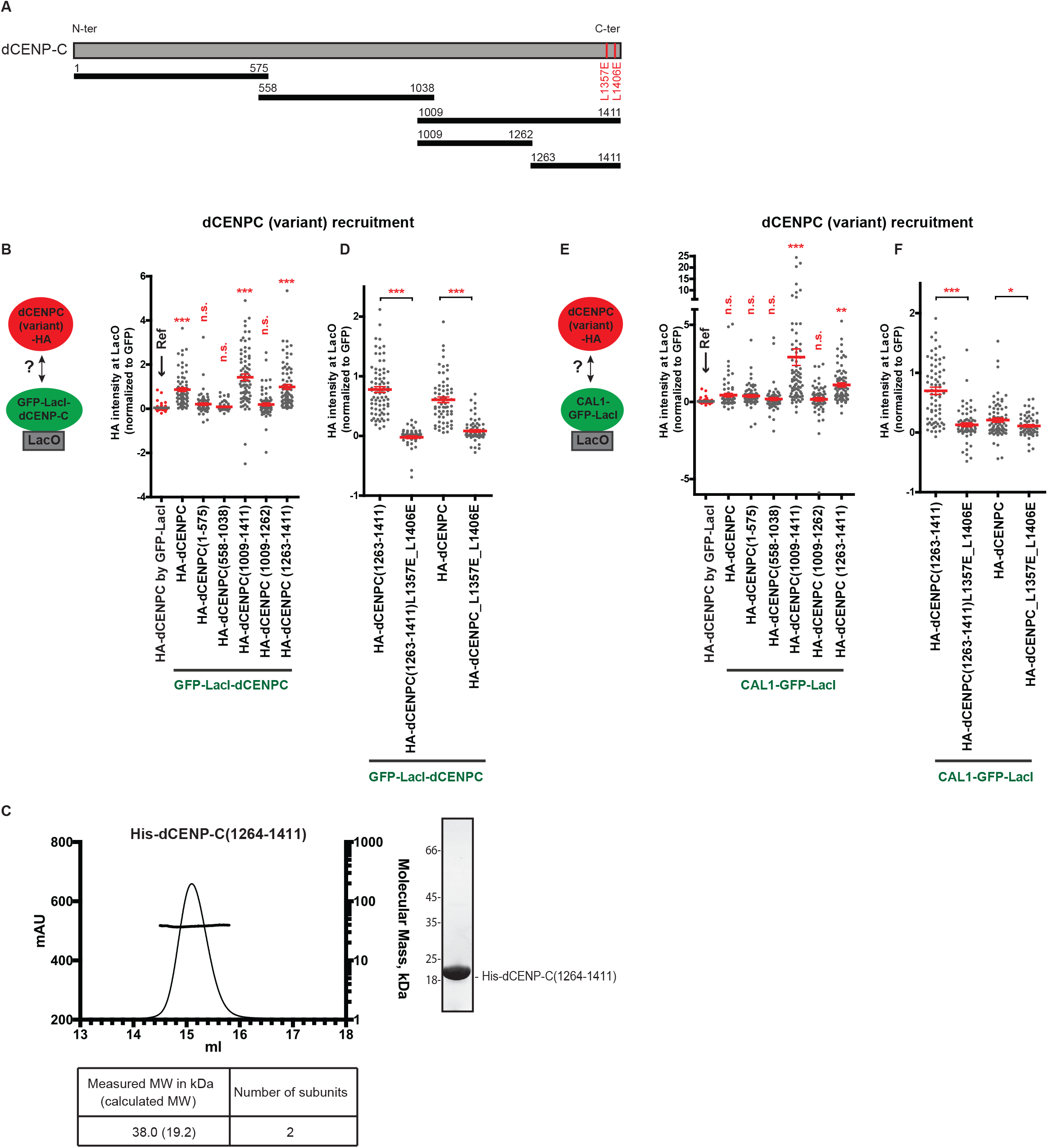
dCENP-C dimerization is necessary for its interaction with CAL1. (A) The dCENP-C N-terminal, middle and C-terminal regions used in this study are indicated by black horizontal lines with numbers indicating amino-acid positions. (B, D) Experimental scheme and quantitation of normalized HA-tagged dCENP-C and dCENP-C fragment mean intensities at LacO upon tethering of GFP-LacI-dCENP-C or GFP-LacI (B, first lane). (C) SEC MALS experiments performed with recombinant His-dCENP-C(1264-1411). Elution volume (ml, x-axis) is plotted against absorption at 280 nm (mAu, left y-axis) and molecular mass (kDa, right y-axis). Tables show measure molecular weight (MW) and calculated MW. Mw/Mn = 1.001. (E, F) Experimental scheme and quantitation of normalized HA-tagged dCENP-C and dCENP-C fragment mean intensities at LacO upon tethering of CAL1-GFP-LacI or GFP-LacI (E, first lane). (See also Figure S6). Error bars show SEM (*P < 0.05; ** P < 0.01; *** P < 0.001; (n.s.) as not significant).

The crystal structure of Mif2p revealed that 3 residues in the cupin-fold are required for Mif2p dimer formation (Cohen et al., 2008). Comparative structure analysis allowed prediction of a putative *Drosophila* CENP-C dimer and identified two Leucine residues (position 1357 and 1406) that could be required for dCENP-C dimer formation (Figures 6A and S6B). Mutations of these two residues (L1357E and L1406E) in the cupin fragment alone (1263-1411) or in full length dCENP-C are sufficient to prevent its dimerization (Figures 6D and S6A), indicating that these two Leucines are crucial for the formation of dCENP-C dimers.

To determine whether the dimerization domain of dCENP-C is required for association with CAL1, we tested the recruitment of the various dCENP-C fragments to the LacO arrays by CAL1-GFP-LacI (Figures 6E and S6C). Our analysis revealed that CAL1 associates with the dCENP-C(1263-1411) fragment, confirming previous observations using yeast two-hybrid analysis (Schittenhelm et al., 2010). Thus, the dCENP-C dimerization domain and the domain of interaction with CAL1 reside within the same region of dCENP-C. Importantly, mutations of the two Leucine residues in the dimerization domain of dCENP-C also prevents its recruitment by LacO-tethered CAL1-GFP-LacI (Figures 6F and S6C). This suggests that dimerization of the cupin-fold of dCENP-C is necessary to associate with CAL1.

### CAL1, dCENP-C and dCENP-A are sufficient for dCENP-A propagation in human cells

To examine if the three *Drosophila* centromere proteins are sufficient to stably propagate dCENP-A in a heterologous system across continuous cell division cycles, we generated a U2OS-LacO line stably expressing CAL1-V5, HA-dCENP-C and dCENP-A (Figure S7A). For comparison, we utilized control cells stably only expressing dCENP-A. Both cell lines were then transiently transfected with GFP-LacI-dCENP-C to initiate dCENP-A loading at the LacO arrays (Figures 7A, 7B and S7B). After 7 days of growth without selection, the GFP-LacI-CENP-C signal was no longer detectable at the LacO. Then a second transfection of mCherry-tagged dCENP-A was performed to assay *de novo* loading, while we utilized TetR-Myc to visualize the arrays (Figures 7A and 7C). Whereas we could never observe the recruitment of dCENP-A-mCherry at LacO arrays in cell lines stably expressing only dCENP-A, new loading of dCENP-A-mCherry occurred at LacO arrays in a small subset of cells stably expressing all three centromere proteins (Figure 7C). Interestingly, we noticed that in the latter cell line, dCENP-A also localizes to endogenous human centromeres (data not shown), but decided for this study to only focus on the inheritance of dCENP-A at the LacO array. We conclude that after temporary targeting dCENP-C to an ectopic chromosomal site in human cells, the *Drosophila* centromere proteins dCENP-C, CAL1 and dCENP-A are sufficient to create a self-propagating epigenetic loop to ensure centromere dCENP-A inheritance from one cell generation to the next.

**Figure 7:**
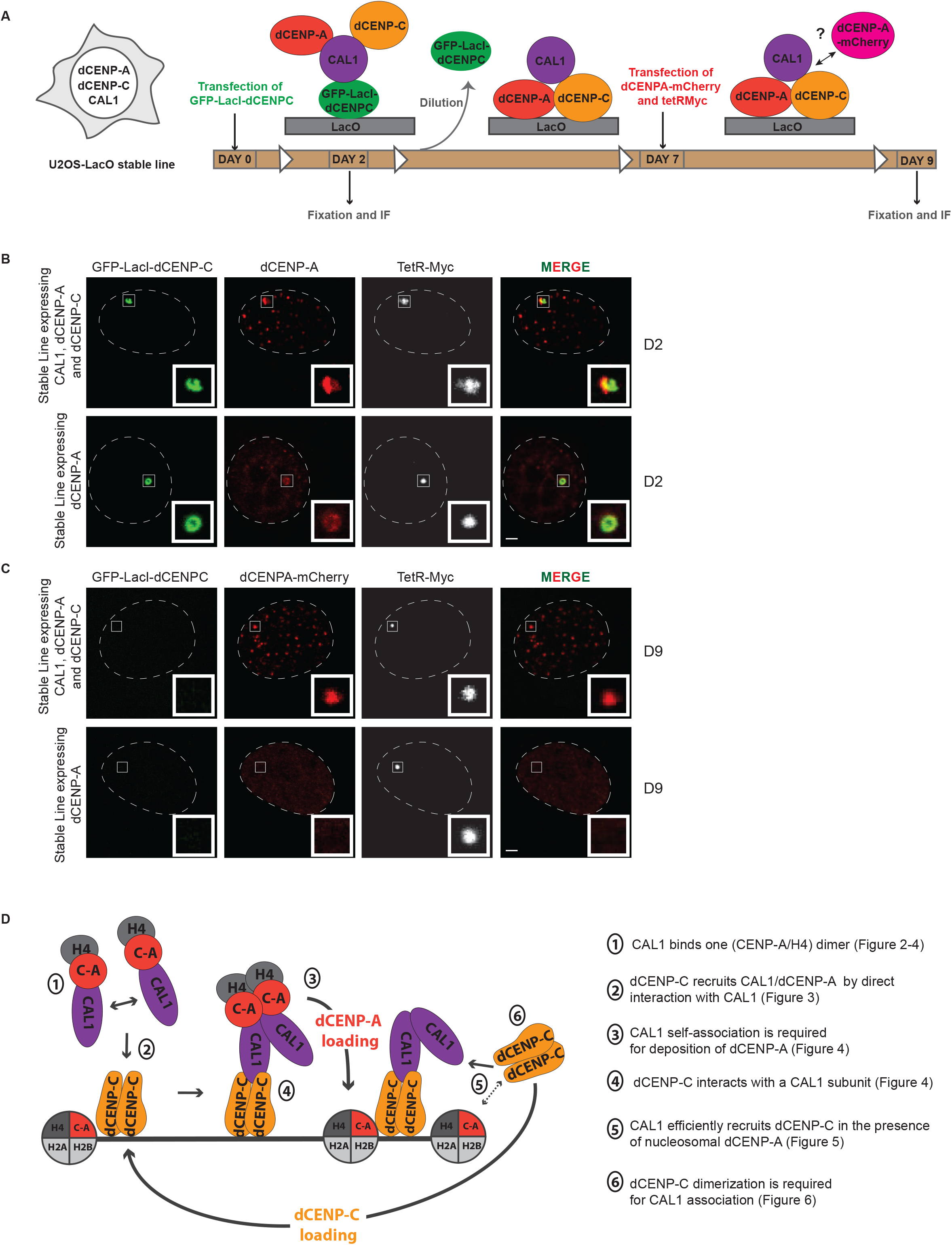
dCENP-A can be stably inherited at ectopic chromosomal sites where *Drosophila* centromere factors are artificially targeted. (A) Experimental procedure for the IF experiments showed in (B, C). (B, C) Representative IF images of the recruitment to the LacO arrays of dCENP-A 2 days (D2) after transfection of GFP-LacI-dCENP-C (B) and dCENP-A-mCherry 9 days (D9) after transfection of GFP-LacI-dCENP-C (C) in U2OS-LacO stable cell line expressing the 3 centromere proteins or only dCENP-A. Insets show magnification of the boxed regions. Scale bar, 5μm. (See also Figure S7). (D) Model for centromere epigenetic inheritance in *Drosophila.* For dCENP-A (C-A in red) loading, a dCENP-C dimer (in yellow) recruits a CAL1 dimer (in purple) in complex with 2x (dCENP-A /H4). For dCENP-C recruitment, one subunit of the CAL1 dimer is available for recruiting a soluble dCENP-C dimer, which is stabilized at the chromatin by dCENP-A nucleosomes.

## DISCUSSION

Understanding how epigenetic marks are propagated through the cell cycle to maintain chromatin identity is a key question in chromosome biology. The development of a heterologous system to investigate centromere inheritance in *Drosophila* was used to dissect step-by-step the roles of the three *Drosophila* centromere components dCENP-A, dCENP-C and CAL1 in this process.

We find that the CATD is partially conserved between *Drosophila* and human as the dCATD is larger and requires the majority of the histone fold domain. As observed for humans, we also show that the dCATD of dCENP-A is similar to the domain recognized by the dCENP-A chaperone CAL1.

Importantly, our findings shed light on the mechanism of dCENP-C recruitment to chromatin, allowing us to close the epigenetic loop and propose a model for centromere propagation (Figure 7D). For dCENP-A loading, dCENP-C plays the role of the recruiting factor: a dCENP-C dimer recruits CAL1, likely in form of a dimer in complex with 2x (dCENP-A/H4). Although the monomeric N-terminal fragment of CAL1 can form a trimeric complex with 1x dCENP-A/H4, we observe that CAL1’s ability to oligomerize is necessary for incorporation of dCENP-A into chromatin, thus suggesting that a dCENP-A/H4 tetramer is required for nucleosome formation. Although plasmid supercoiling assay previously demonstrated that the subregion (1-160) of CAL1 lacking the self-association domain is sufficient for dCENP-A nucleosome assembly activity *in vitro* (Chen et al., 2014), excess of dCENP-A/H4/CAL1 complex under these conditions might be sufficient to overcome the self-association requirement observed *in vivo.* A role for dCENP-C as a stable platform for CAL1/dCENP-A recruitment is further supported by experiments indicating that dCENP-C is stably bound to centromeres (Lidsky et al., 2013; Mellone et al., 2011).

However, dCENP-C recruitment and maintenance at centromeres remain poorly understood. Here, we demonstrate that LacO-tethered CAL1 efficiently recruits dCENP-C in the presence of dCENP-A. Interestingly, although dCENP-C dimerization is required to interact with CAL1, a non-self-associating, monomeric fragment of CAL1 is sufficient to bind a dCENP-C dimer. This finding supports an interesting model, in which one subunit of a potentially CAL1 dimer interacts with a chromatin-bound dCENP-C dimer, while the other CAL1 subunit is available for recruiting a new soluble dCENP-C dimer (Figure 7D). CAL1 is unlikely to be required for dCENP-C maintenance, as in contrast to dCENP-C, it was shown to be very dynamic at centromeres (Lidsky et al., 2013; Mellone et al., 2011). In addition, we demonstrate that CAL1 does not bind nucleosomal dCENP-A, suggesting that CAL1 dissociates from the chromatin after dCENP-A loading. Significant enhancement of dCENP-C localization at the LacO by the presence of dCENP-A nucleosomes instead suggests that dCENP-C maintenance might involve interaction with chromatin bound dCENP-A, similar to the human situation. This model also explains why dCENP-C was found to be dependent on both CAL1 and dCENP-A for its centromere targeting (Erhardt et al., 2008; Goshima et al., 2007; Schittenhelm et al., 2010). In agreement, we provide evidence for direct, albeit weak interactions between dCENP-C and dCENP-A. Our finding that nucleosomal dCENP-A is required for dCENP-C recruitment suggests that the direct interactions between dCENP-C and dCENP-A occur most robustly on assembled dCENP-A chromatin. In addition to direct interaction with dCENP-A, insensivity of dCENP-C to IPTG treatment after LacI recruitment could also involve its affinity to DNA. Indeed, it was reported in human, rat and yeast that CENP-C has DNA binding property (Cohen et al., 2008; Kato et al., 2013; Sugimoto et al., 1994). CAL1 contains properties of both HJURP and the Mis18 complex through its ability to act as the chaperone for dCENP-A and bind centromeric dCENP-C, respectively. In addition, CAL1 also functions in the recruitment of dCENP-C to the centromere and acts as a potential loading factor. Interestingly its human functional homolog HJURP was also shown to recruit a C-terminal fragment of hCENP-C, although recruitment of full-length CENP-C could not be detected (Tachiwana et al., 2015). So, despite the poor conservation between the centromere factors of *Drosophila* and humans, our study reveals remarkable similarities in their behavior, suggesting a general conserved mechanism for centromere inheritance in these organisms.

In summary, we demonstrate that an epigenetic loop of dCENP-A loading and self-propagation can be ‘kick-started’ by transiently targeting dCENP-C to the ectopic LacO regions in a heterologous system, identifying the centromere factors dCENP-A, dCENP-C and CAL1 as sufficient for dCENP-A inheritance.

## EXPERIMENTAL PROCEDURES

### Cell culture and transfection

U2OS cells containing 200 copies of an array of 256 tandem repeats of the 17 bp LacO sequence on chromosome 1 (gift from B.E Black, University of Pennsylvania, Philadelphia; Janicki et al. 2004) were cultured in DMEM supplemented with 10% FBS, 100 U/ml penicillin, 100 μg/ml streptomycin and selected with 100 μg/ml Hygromycine B at 37°C in a humidified incubator with 5% CO2. Schneider S2 cells were grown at 25°C in Schneider’s Drosophila medium (Serva) supplemented with 10% FBS (Sigma-Aldrich) and antibiotics (300 μg/ml penicillin, 300 μg/ml streptomycin, and 750 μg/ml amphotericin B).

U2OS cells were transfected with FuGENE HD Transfection Reagent (Promega) and Schneider S2 cells with XtremeGENE DNA transfection reagent (Roche).

### Plasmids

All *Drosophila* expression vectors used in this study were constructed in a pMT/V5-6his vector or a pDS47 vector. All mammalian expression vectors used in this study were constructed in a pN2-CMV vector.

#### Drosophila expression vectors

| **dCENP-A/H3 chimera vectors** | **CAL1 vectors** |
| --- | --- |
| pDS47_dCENP-A_HA<br>pDS47_H3_HA<br>pDS47_H3<sup>L1α2</sup>_HA<br>pDS47_dCENP-A<sup>C</sup>_HA<br>pDS47_dCENP-A<sup>α2+C</sup>_HA<br>pDS47_H3<sup>L1α2+C</sup>_HA<br>pDS47_H3<sup>α2+C</sup>_HA<br>pDS47_H3<sup>C</sup>_HA<br>pDS47_dCENP-A<sup>L1α2</sup>_HA<br>pDS47_dCENP-A<sup>L1α2+C</sup>_HA<br>pDS47_H3<sup>L1+C</sup>_HA<br>pDS47_H3<sup>L1α2+C(223)</sup>_HA<br>pDS47_H3<sup>L1α2+C(222)</sup>_HA | pMT_CAL1_HA<br>pMT_CAL1(NT)_HA<br>pMT_CAL1(CT)_HA |
| pDS47_H3<sup>L1α2+C(219)</sup>_HA<br>pDS47_H3<sup>L1α2+C(218)</sup>_HA<br>pDS47_H3<sup>L1α2+C(206)</sup>_HA |  |

#### Mammalian expression vectors

*Mammalian expression vectors:*
| **GFP-LacI-tagged vectors** | **V5-tagged vectors** |
| --- | --- |
| pN2_CMV_dCENP-A_GFPLacI<br>pN2_CMV_H3_GFPLacI<br>pN2_CMV_H3<sup>L1α2</sup>_GFPLacI<br>pN2_CMV_dCENP-A<sup>C</sup>_GFPLacI<br>pN2_CMV_dCENP-A<sup>α2+C</sup>_GFPLacI<br>pN2_CMV_H3<sup>L1α2+C</sup>_GFPLacI<br>pN2_CMV_H3<sup>α2+C</sup>_GFPLacI | pN2_CMV_CAL1_V5<br>pN2_CMV_dCENP-A_V5<br>pN2_CMV_tetR_V5<br>pN2_CMV_dCENP-A<sup>C</sup>_V5<br>pN2_CMV_dCENP-A<sup>C</sup>_V5 |
|  | **HA-tagged vectors** |
| pN2_CMV_H3<sup>C</sup>_GFPLacI<br>pN2_CMV_dCENP-A<sup>L1α2</sup>_GFPLacI<br>pN2_CMV_dCENP-A<sup>L1α2+C</sup>_GFPLacI<br>pN2_CMV_H3<sup>L1+C</sup>_GFPLacI<br>pN2_CMV_GFPLacI_dCENP-C(CT)<br>pN2_CMV_GFPLacI<br>pN2_CMV_GFPLacI_HJURP<br>pN2_CMV_CAL1_GFPLacI<br>pN2_CMV_GFPLacI_dCENP-C | pN2_CMV_HA_dCENP-C<br>pN2_CMV_CAL1_HA<br>pN2_CMV_dCENP-A_HA<br>pN2_CMV_H3_HA<br>pN2_CMV__CAL1(NT)_HA<br>pN2_CMV__CAL1(CT)_HA<br>pN2_CMV__CAL1(MID)_HA<br>pN2_CMV__CAL1(1-160)_HA<br>pN2_CMV__CAL1(101-407)_HA<br>pN2_CMV_HA_dCENP-C(NT)<br>pN2_CMV_HA_dCENP-C(CT)<br>pN2_CMV_HA_dCENP-C(MID)<br>pN2_CMV_HA_dCENP-C(1009-1262)<br>pN2_CMV_HA_dCENP-C(1263-1411)<br>pN2_CMV_HA_dCENP-C(1263-1411)L1357E-L1406E<br>pN2_CMV_HA_dCENP-C_L1357E-L1406E |
| ***GFP-LacIw-tagged vectors*** |  |
| pN2_CMV_CAL1(NT)_GFPLacIw<br>pN2_CMV_CAL1(1-160)_GFPLacIw<br>pN2_CMV_GFPLacIw<br>pN2_CMV_dCENP-A(Y187A-L188Q)_GFPLacIw<br>pN2_CMV_dCENP-A<sup>C</sup>_GFPLacIw<br>pN2_CMV_dCENP-A_GFPLacIw<br>pN2_CMV_CAL1_GFPLacIw<br>pN2_CMV_GFPLacIw_dCENP-C |  |
| ***Myc- and mCherry-tagged vectors*** | ***GoldenGate vectors*** |
| pN2_CMV_tetR_Myc<br>pN2_CMV_dCENP-A_mCherry | pN2_CMV_dCENP-A_P2A_HAdCENP-C<br>pN2_CMV_CAL1V5_T2A_dCENP-A<br>pN2_CMV_CAL1V5_T2A_dCENP-A_T2A_HAdCENP-C |

As originally described (Straight et al., 1996), the LacI protein used in this study contained an 11-amino-acid-C-terminal truncation to prevent tetramer formation and to avoid artefactual linkage of daughter chromatids during mitosis when targeted to the chromosomally-integrated LacO array. This LacI protein is insensitive to IPTG treatment. Therefore, for all experiments in which the LacI fusion needed to be displaced from the LacO array, a double mutant version of LacI, termed “LacI weak” was used. This mutant was derived from a construct obtained from Taddei and colleagues (Loiodice et al., 2014) and comprises three mutations towards the N-terminus of the protein (G58D, Q60GGG and T68S) in addition to the C-terminal truncation.

For the cloning of vectors with multiple genes under the same promoter, we used the Golden Gate assembly method as described in (Engler and Marillonnet, 2014; Szymczak et al., 2004) and the genes were separated with T2A or P2A sequences to create a self-cleaving peptide.

For expression of the recombinant proteins in SEC-MALS and gel filtration assays, dCENP-A and dH4 were cloned into a pET3a vector, CAL1(1-160) into a pEC-K-3C-His-GST vector, dCENP-C and hH4 into a pEC-K-3C-His vector, all containing a T7 promoter.

#### Immunoflorescence

Cells were harvested 48 h (for U2OS-LacO cells) or 72 h (for S2 cells) after transfection and fixed for 10 min in 3.7% formaldehyde in PBS and 0.1% Triton X-100. Primary antibodies were incubated overnight at 4°C at the following dilutions or concentrations: rat anti-HA (1:20; clone 3F10; E. Kremmer), rabbit anti-hCENP-C (1:100; B. Black; Bassett et al., 2010), mouse anti-hCENP-A (Abcam, ab139139), rabbit anti-dCENP-C (1:100; a gift from C. Lehner, University of Zurich, Zurich, Switzerland; Heeger et al., 2005), rabbit anti-V5 (1:500; Sigma V8137), mouse anti-V5 (1:100; Invitrogen, R96025), rabbit anti-Myc (1:100; Abcam ab9106), chicken anti-dCENP-A (1:100), mouse anti-mCherry (1:15; kindly provided by S. Sacchani). Goat secondary antibodies conjugated to Alexa Fluor 488, Alexa Fluor 555 or Alexa Fluor 647 (Invitrogen) were incubated for 1 h at room temperature at a dilution of 1:100. DNA was counterstained with DAPI at 5 μg/ml.

For experiments involving IPTG treatment on U20S-LacO cells, cells were treated 48 h after transfection with 15 mM IPTG (Sigma-Aldrich) for 1h and processed for immuno-fluorescence, as described earlier in this section.

#### Microscopy and image analysis

Two microscopes were used to image the samples. With a microscope DeltaVision RT Elite, images were acquired as 45-50 z stacks of 0.2 μm increments using a 100× oil immersion objective and a monochrome camera (CoolSNAP HQ; Photometrics) and deconvolved using softWoRx Explorer Suite (Applied Precision). With an Olympus confocal microscope FV1200, images were acquired as 10-20 z stacks of 0.42μm increments using a 60× oil immersion objective.

Quantification of fluorescence intensities was performed using ImageJ. For experiments with S2 cells, the mean fluorescence intensity of the protein of interest was measured at centromeres and subtracted from the average of the mean fluorescence intensities of three points randomly chosen in the nucleus (background), then normalized to the mean fluorescent intensity of centromeres (marked with anti dCENP-C or dCENP-A antibodies). For the experiments with U2OS-LacO cells, the mean fluorescence intensity of the protein of interest was measured at the LacO spot, subtracted from the mean fluorescence intensity in the nucleus (background) and normalized to the GFP (in case of GFP-LacI tagged proteins), V5 or Myc (tetR-V5 or tetR-Myc in experiments with IPTG treatment) mean fluorescence intensity of the corresponding LacO spot. A minimum of 20 cells were analysed per biological replicate, and a minimum of two independent biological replicates were quantified per experiment. GraphPad Prism software was used for statistical analysis and graphical representations. Unpaired student T-tests were performed in Figures 1D, 3B, 3C, 3H, 3I, 5B, 5D, 6C, 6F, S5A and S5B; one-way ANOVA with Dunnett’s post-test in Figures 1C, 2B, 4B, 4C, 4H, 5E, 6B, 6E and S4D; and oneway ANOVA with Turkey’s post-test in Figures S1C, 1D and S4B. Error bars represent mean +/− SEM.

## SEC MALS

Size-exclusion chromatography (ÄKTAMicro^™^, GE Healthcare) coupled to UV, static light scattering, and refractive index detection (Viscotek SEC-MALS 20 and Viscotek RI Detector VE3580; Malvern Instruments) was used to determine the absolute molecular mass of proteins and protein complexes in solution. Injections of 100 μl of 1-4 mg/ml material were run on a Superdex 200 10/300 Increase GL (GE Healthcare) size-exclusion column pre-equilibrated in 50 mM HEPES (pH 8.0), 150 or 300 mM NaCl, and 1 mM TCEP at 22°C with a flow rate of 0.5 ml/min. Light scattering, refractive index (RI), and A280 _nm_ were analyzed by a homo-polymer model (OmniSEC software, v5.02; Malvern Instruments) using the following para-meters for: ∂A280nm/∂c = 1.04 AU ml mg ^1^ (Cal1 1-160), 0.70 AU ml mg^1^ (Cal1 1-160/H4/CID 101-end), 0.79 AU ml mg ^1^ (Cal1 1-160/H4/CID 144-end), 0.75 AU ml mg ^1^ (His CENP-C 1264-end), ∂n/∂c = 0.185 ml g ^1^ and buffer RI value of 1.335. The mean standard error in the mass accuracy determined for a range of protein–protein complexes spanning the mass range of 6–600 kDa is ±1.9%.

## AUTHOR CONTRIBUTIONS

P.H. and V.R. conceived the project and wrote the paper. V.R. and B.M-P. performed the experiments. All authors designed the experiments. J.A. provided expertise and feedback.

## ACKNOWLEDGMENTS

We thank Ben Black, Susan Janicki, Simona Sacchani, Christian Lehner, Stefan Heidmann for the human U2OS-LacO cells, antibodies and reagents. We also thank Robin Allshire and the Heun lab for helpful comments on the manuscript and Martin Wear, Daniela Venegas, Vasiliki Lazou and Thomas van Emden for technical assistance. We are thankful to Maria Dominguez Castellano for her hosting support. V.R. and P.H. were supported by the Alexander Von Humboldt Foundation, the European Research Council Starting-Consolidator Grant 311674-BioSynCen and a Wellcome Trust Senior Fellowship award. A.A.J. and B.M. are funded by the Wellcome Trust through a Wellcome Research Career Development (095822) and a Senior Research (202811) Fellowships and by the European Commission Marie-Curie Career Integration Grant (334291). The Wellcome Trust Centre for Cell Biology is supported by core funding from the Wellcome Trust (092076).

## SUPPLEMENTAL INFORMATION

**Figure S1:**
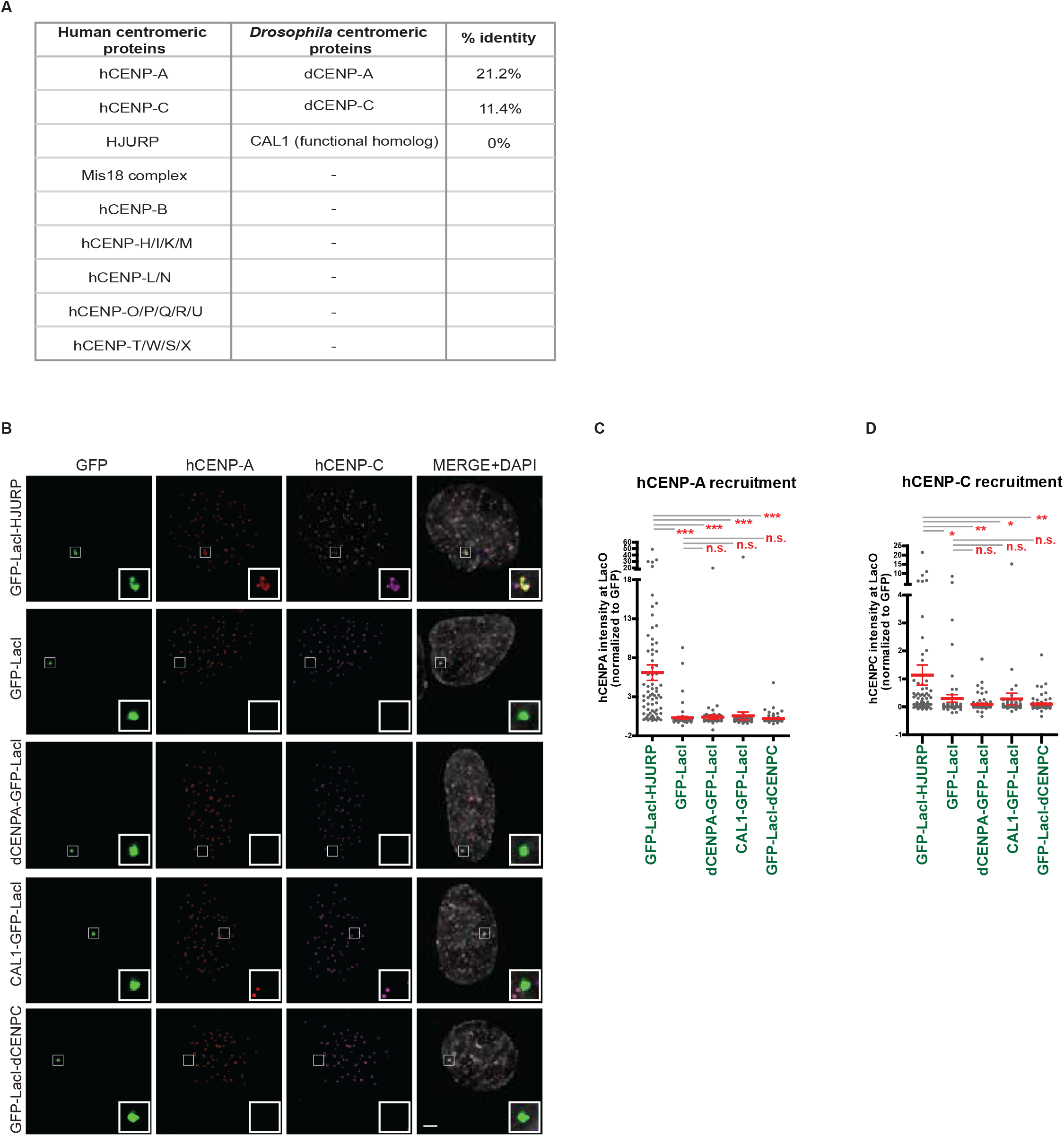
There is no interference of the endogenous human centromere system with the exogenous *Drosophila* centromere system. (A) Table showing centromere protein homologies between Human and *Drosophila.* (B) Representative IF images of hCENP-A and hCENP-C recruitment to the LacO arrays by the *Drosophila* centromere factors fused to GFP-LacI, HJURP-GFP-LacI and the control GFP-LacI in U2OS cells. (C, D) Quantitation of normalized hCENP-A (C) and hCENP-C (D) mean intensities at LacO upon tethering of the indicated proteins fused to GFP-LacI. Insets show magnification of the boxed regions. Scale bar, 5μm. Error bars show SEM (*P < 0.05; ** P < 0.01; *** P < 0.001; (n.s.) as not significant).

**Figure S2:**
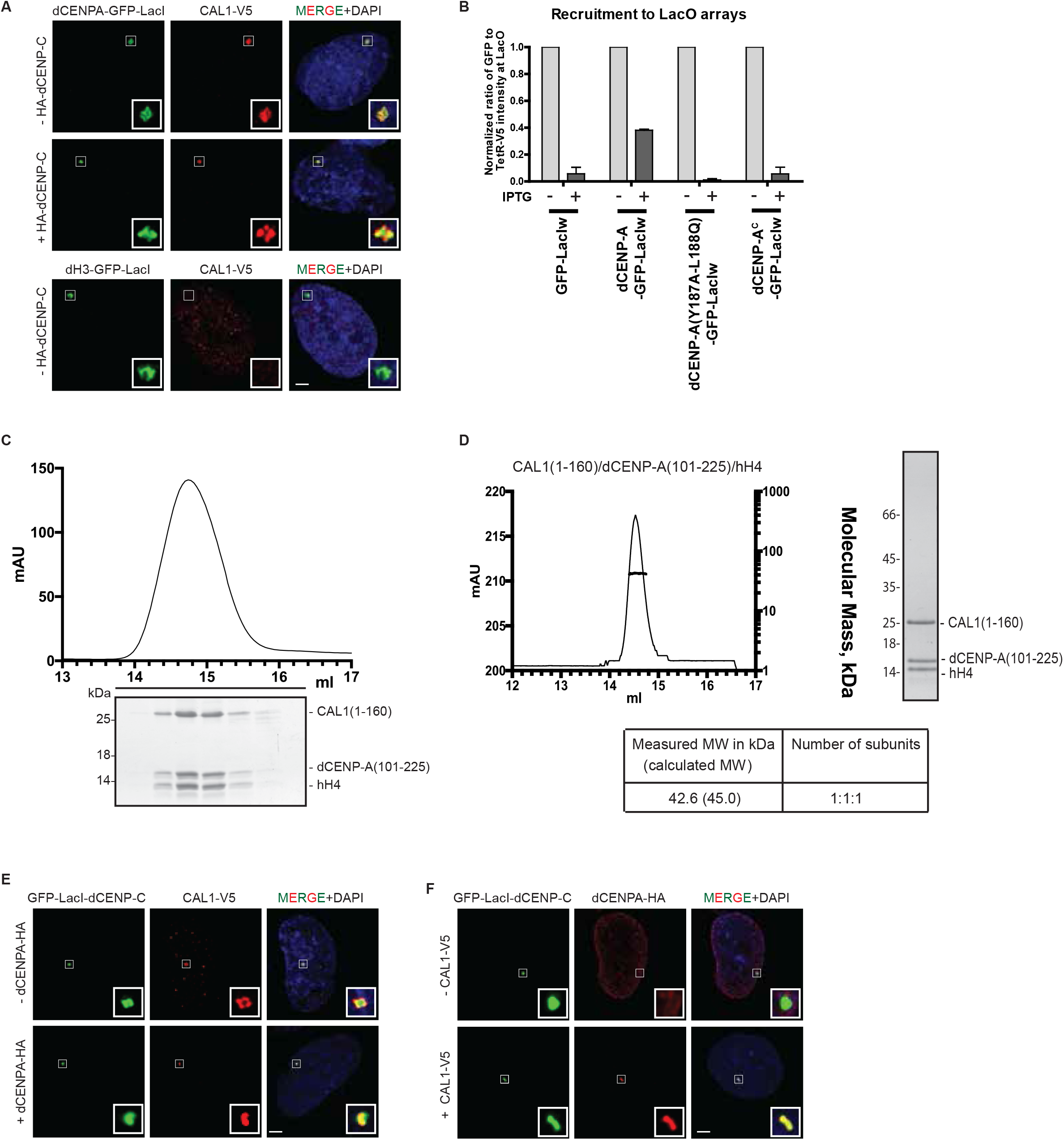
dCENP-C is the factor recruiting the complex CAL1/dCENP-A to the chromatin. (A) Representative IF images of CAL1-V5 recruitment to the LacO arrays by GFP-LacI-tagged dCENP-A and dH3, +/− HA-dCENP-C, in U2OS cells. (B) Quantitation of normalized mean intensities at LacO of the indicated dCENP-A WT and chimeras fused to GFP-LacIw, +/− IPTG treatment. (C) SEC profiles and respective SDS-PAGE analysis of CAL1(1-160) in complex with dCENP-A(101-225) and hH4 complex separated on a S200 increase 10/300. (D) SEC-MALS experiments performed with recombinant CAL1 (1-160) in complex with dCENP-A(101-225) and hH4. Elution volume (ml, x-axis) is plotted against absorption at 280 nm (mAu, left y-axis) and molecular mass (kDa, right y-axis). Tables show measure molecular weight (MW) and calculated MW. Mw/Mn = 1.001. (E, F) Representative IF images of the recruitment to the LacO arrays by GFP-LacI-dCENP-C of CAL1-V5, +/− dCENP-A-HA (E), and dCENP-A-HA, +/− CAL1-V5 (F), in U2OS cells. Insets show magnification of the boxed regions. Scale bar, 5μm.

**Figure S3:**
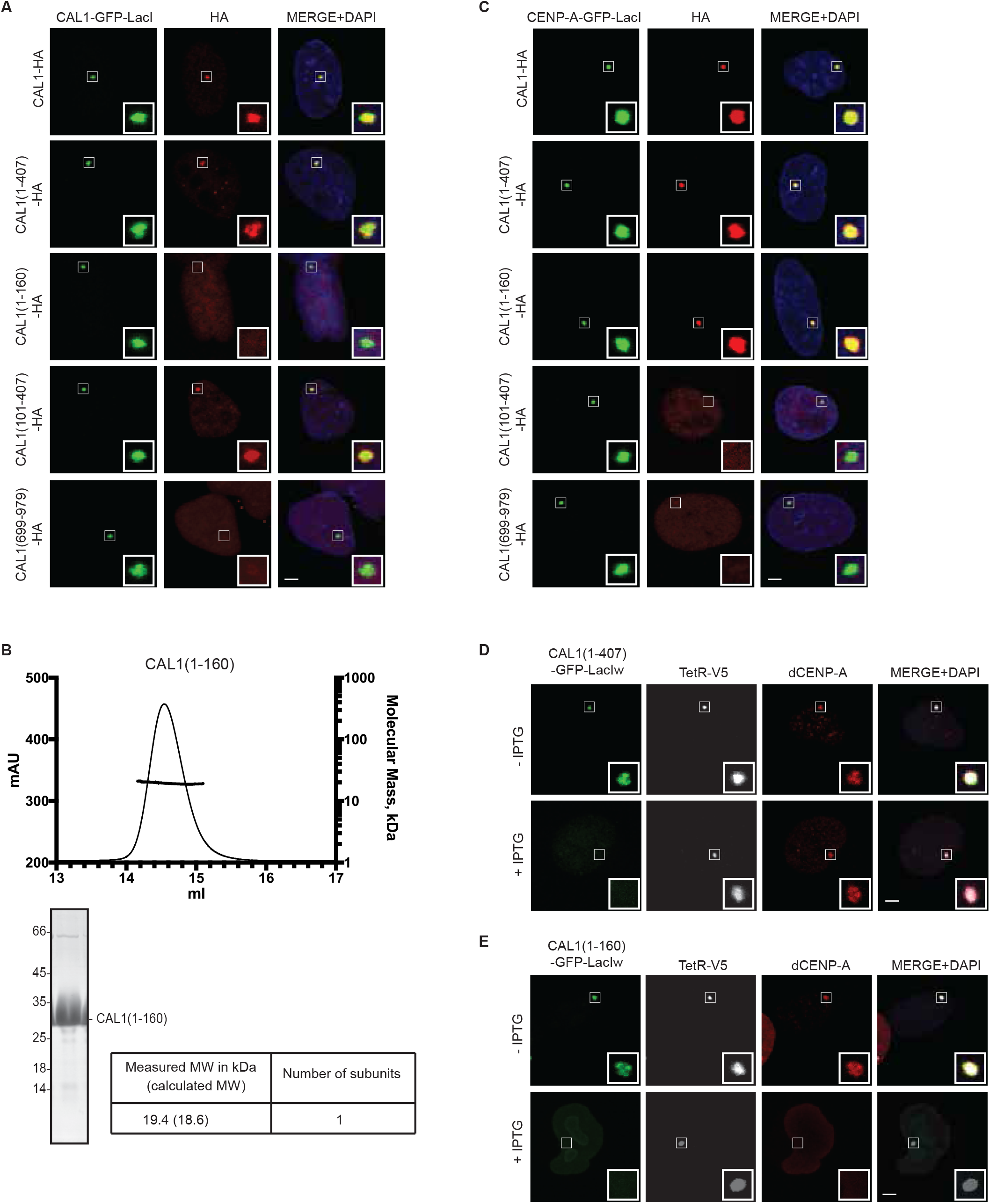
CAL1 self-association is not required for binding but for deposition of dCENP-A. (A, C) Representative IF images of HA-tagged CAL1 and CAL1 fragment recruitment to the LacO arrays by CAL1-GFP-LacI (A) and dCENP-A-GFP-LacI (C) in U2OS cells. (B) SEC MALS experiments performed with recombinant CAL1(1-160). Elution volume (ml, x-axis) is plotted against absorption at 280 nm (mAu, left y-axis) and molecular mass (kDa, right y-axis). Tables show measure molecular weight (MW) and calculated MW. Mw/Mn = 1.001. (D, E) Representative IF images of dCENP-A recruitment to the LacO arrays by CAL1(1-407) (D) and CAL1(1-160) (E) fused to GFP-LacIw, +/− IPTG treatment in U2OS cells. Insets show magnification of the boxed regions. Scale bar, 5μm.

**Figure S4:**
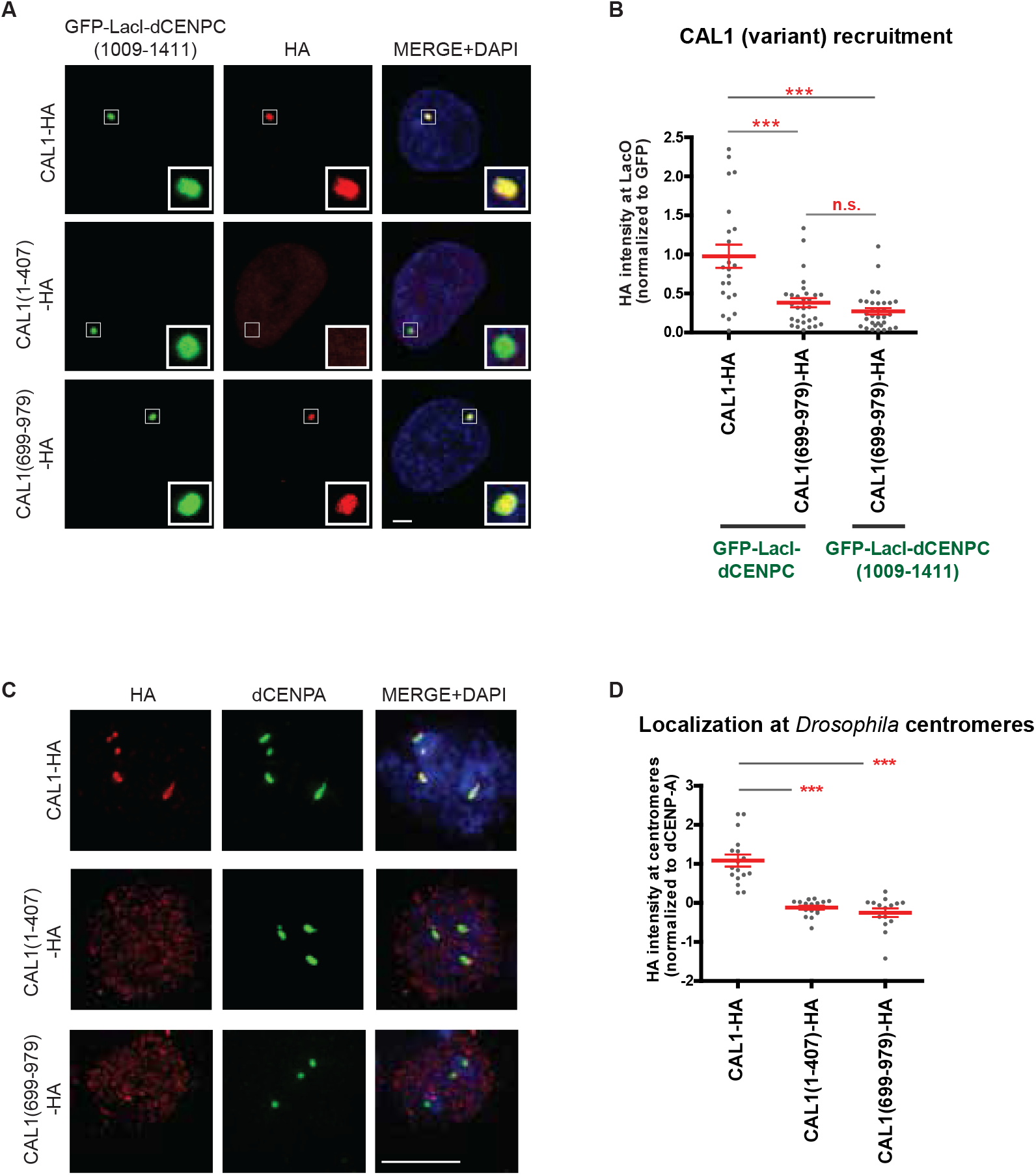
Self-association of CAL1 is required for centromere targeting but not for interaction with dCENP-C. (A) Representative IF images of HA-tagged CAL1 and CAL1 fragment recruitment to the LacO arrays by GFP-LacI-dCENP-C(1009-1411) in U2OS cells. (B) Quantitation of normalized HA-tagged CAL1 and CAL1(699-979) mean intensities at LacO upon tethering of GFP-LacI-tagged dCENP-C and dCENP-C (1009-1411). (C) Representative IF images of HA-tagged CAL1 and CAL1 fragment expression patterns in S2 *Drosophila* cells. dCENP-A marks *Drosophila* centromeres. (D) Quantitation of normalized mean intensities of the indicated CAL1 and CAL1 fragments at centromeres. Insets show magnification of the boxed regions. Scale bar, 5μm. Error bars show SEM (*** P < 0.001; (n.s.) as not significant).

**Figure S5:**
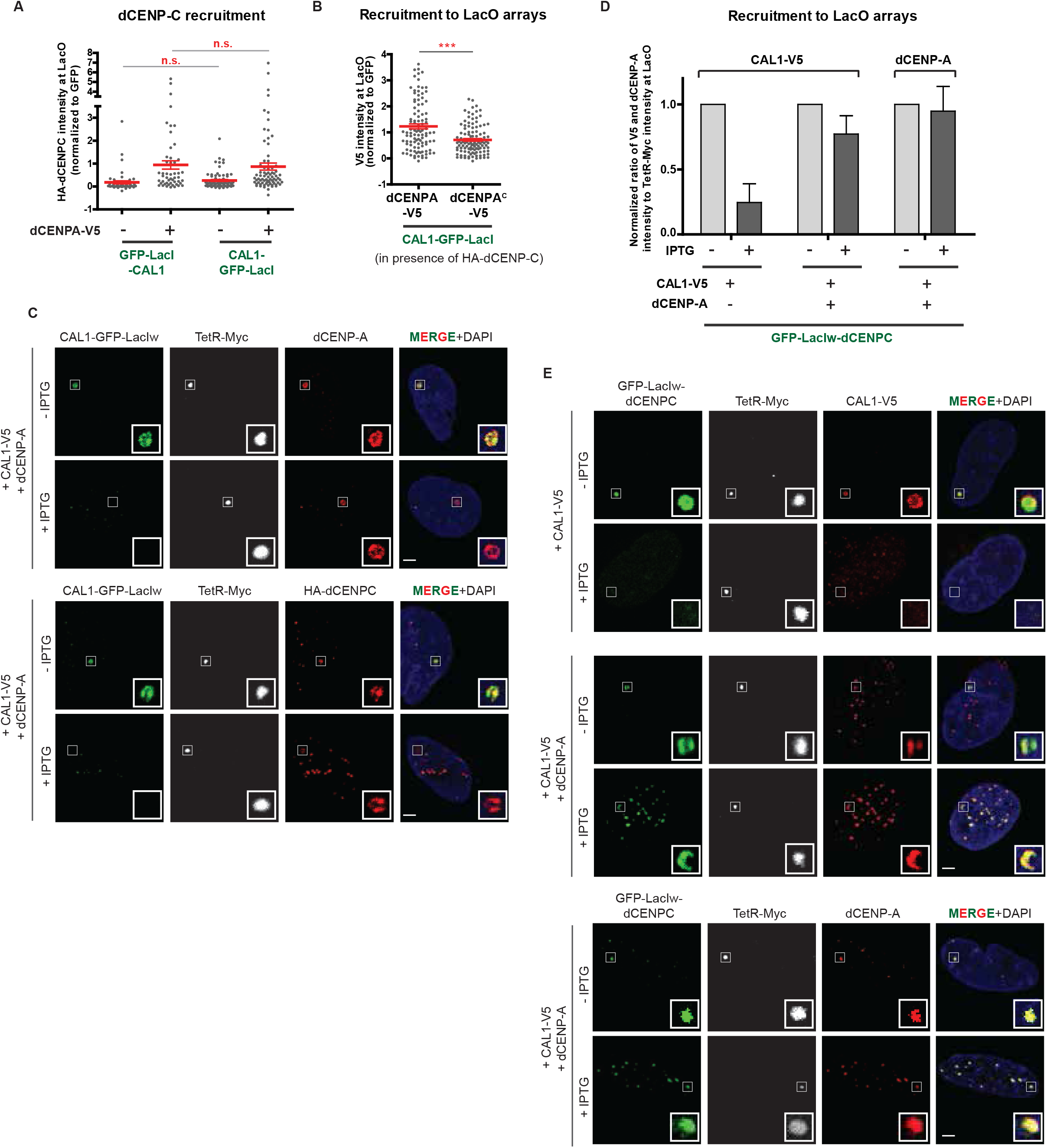
dCENP-C is loaded by CAL1 to the lacO arrays containing dCENP-A nucleosomes. (A) Quantitation of normalized HA-dCENP-C mean intensity at LacO upon tethering of CAL1 fused to GFP-LacI on its N-terminus (GFP-LacI-CAL1) or C-terminus (CAL1-GFP-LacI), +/− dCENP-A-V5. (B) Quantitation of normalized V5-tagged dCENP-A and dCENP-A mean intensities at LacO upon tethering of CAL1-GFP-LacI in presence of HA-dCENP-C. (C) Representative IF images of dCENP-A and HA-dCENP-C recruitment to the LacO arrays by CAL1-GFP-LacIw, +/− IPTG treatment in U2OS cells. (D) Quantitation of normalized CAL1-V5 and dCENP-A mean intensities at LacO upon tethering of GFP-LacIw-dCENP-C, +/−dCENP-A, +/− IPTG treatment. (E) Representative IF images of CAL1-V5 and dCENP-A recruitment to the LacO arrays by GFP-LacIw-dCENP-C, +/−dCENP-A, +/− IPTG treatment in U2OS cells. Insets show magnification of the boxed regions. Scale bar, 5μm. Error bars show SEM (*** P < 0.001; (n.s.) as not significant).

**Figure S6:**
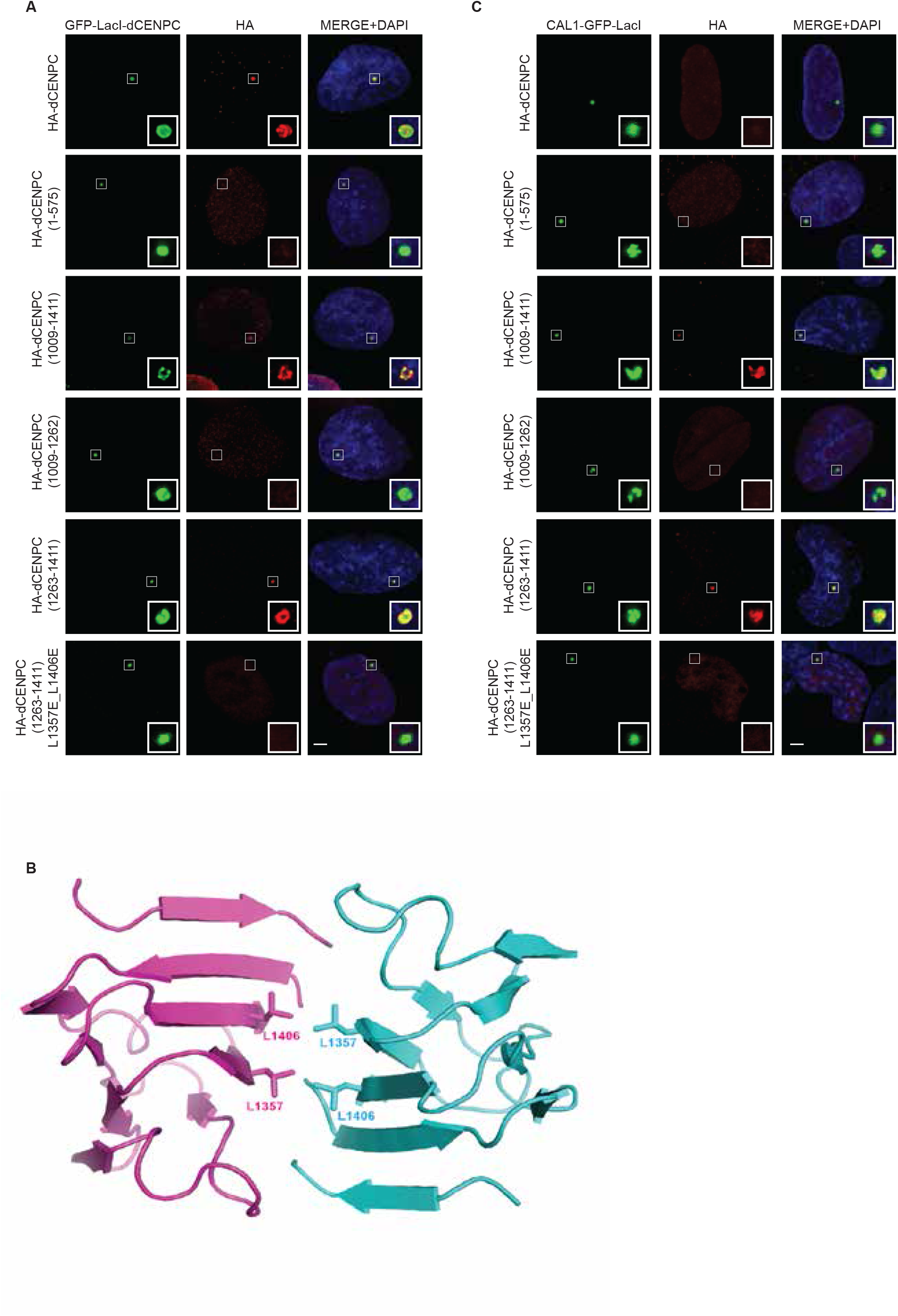
dCENP-C dimerization is necessary for its interaction with CAL1. (A, C) Representative IF images of HA-tagged dCENP-C and dCENP-C fragment recruitment to the LacO arrays by GFP-LacI-dCENP-C (A) and CAL1-GFP-LacI (C) in U2OS cells. (B) Predicted crystal structure of dCENP-C(1302-1409) using Phyre2 (www.sbg.bio.ic.ac.uk/~phyre2/) (Jefferys et al., 2010; Soding, 2005).

**Figure S7:**
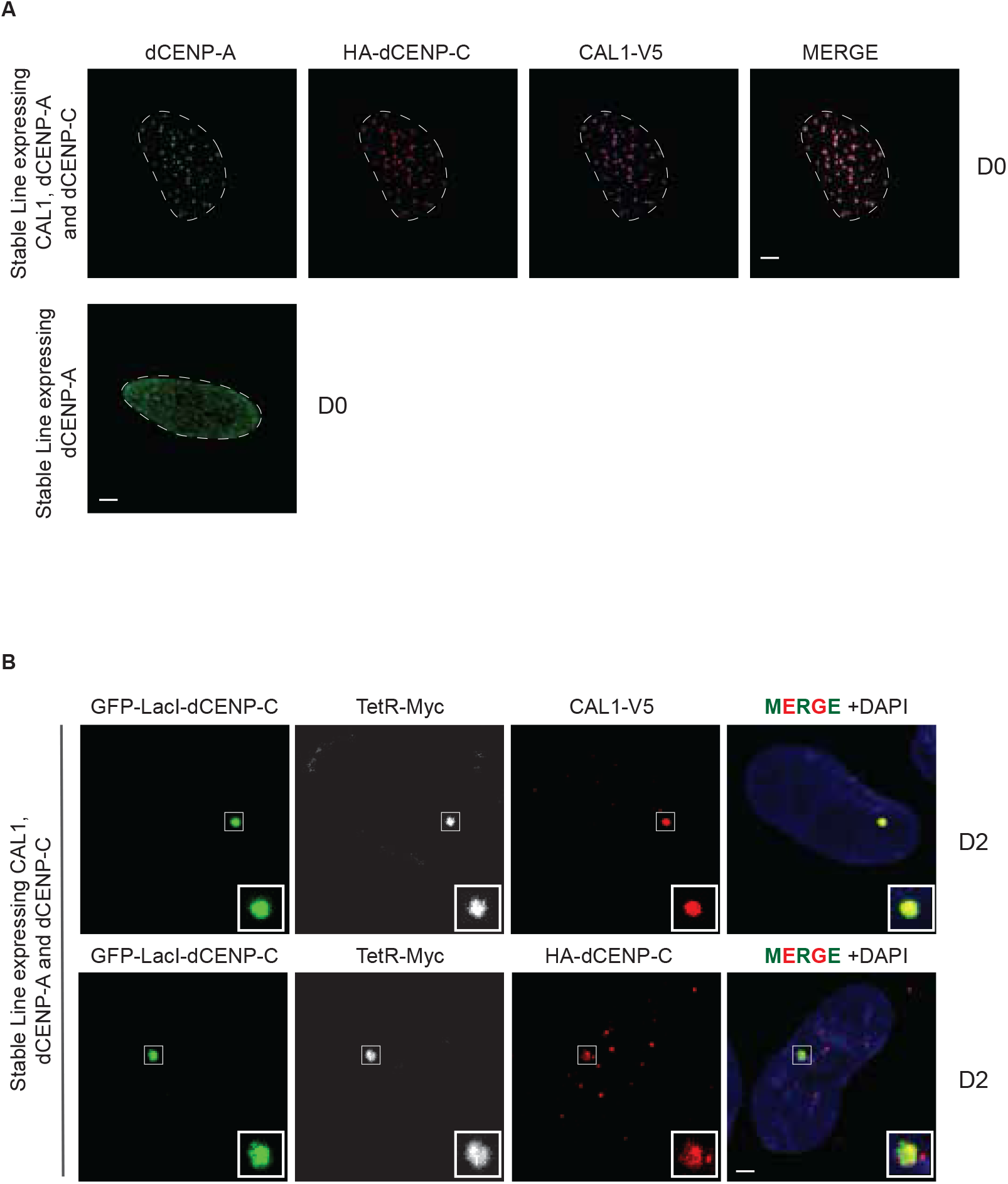
dCENP-A can be stably inherited at ectopic chromosomal sites where Drosophila centromere factors are artificially targeted. (A) Expression of CAL1-V5, dCENP-A and HA-dCENP-C in U2OS-LacO stable cell line expressing the 3 centromere proteins or only dCENP-A (Day 0). (B) Representative IF images of CAL1-V5 and HA-dCENP-C recruitment to the LacO arrays 2 days (D2) after transfection of GFP-LacI-dCENP-C in U2OS-LacO stable cell line expressing the 3 centromere proteins. Insets show magnification of the boxed regions. Scale bar, 5μm.

